# Chronic HDACi emends microglial differentiation and neurological disease in a mouse model of intellectual disability

**DOI:** 10.1101/2025.05.01.651719

**Authors:** Md. Suhail Alam, Prasad Kottayil Padmanabhan, Ryan McArdle, Alejandro Lopez-Ramirez, Arpitha MysoreRajashekara, James Knopp, Gabriela Kim, Maria Virginia Centeo, Apkar Vania Apkarian, Vivek Swarup, Kasturi Haldar

## Abstract

Defect in lysine-specific methyltransferase 2D (KMT2D) underlies a rare intellectual disability disorder, Kabuki Syndrome (KS). We show that in addition to reduction in post-natal neurogenesis, KS mice present small, arrested hippocampal microglia which single cell RNA sequence analyses revealed as globally downregulated, occupying neither activated nor surveillance states. Weekly administration of a triple combination formulation (TCF) containing the FDA-approved HDAC inhibitor Vorinostat, HPBCD and PEG-400 for three months, boosted microglia. Transcriptomic, pseudotime cell trajectory analyses, imaging and behavioral data suggest that TCF differentiated KS microglia from arrested to non-diseased and non-activated surveillance forms by selective acetylation of histone H3, that restored the open chromatin mark H3K4me3 independent of Kmt2D, to improve cognitive and nociceptive responses. Wild-type microglia showed no substantial transcriptional or histone changes, intimating normal heterochromatin resisted TCF. These findings reveal a novel chromatin-mediated mechanism of microglial differentiation uncoupled from activation, with therapeutic potential for KS and related disorders.

## Introduction

Histone deacetylases (HDACs) are key players in epigenetic regulation. Their inhibitors (HDACi) show direct anti-tumor effects as well as target the immune microenvironment that support tumors ^1–3^. HDACi elicit complex cellular responses by blocking multiple classes of HDAC enzymes to induce acetylation of histones and promote the opening of chromatin. Since epigenetic mechanisms are important for brain function at all stages ^4, 5^, HDACi are of interest for their potential to promote many aspects of cerebral function ^6–9^. HDAC-stimulation of post-natal adult neurogenesis ^10^ is expected to improve hippocampal functions of learning and memory. Inhibition/reduction of specific HDACs (like HDAC3) facilitates the microglial response to inflammation^11–13^. However, HDAC functions are essential for the development of the brain ^14–17^. To mitigate, HDAC class-selective inhibitors and HDACi designed to have better brain penetration are emerging ^18–21^. But for presently available FDA-approved HDACi, toxicity and poor blood-brain barrier (BBB) permeability limit dose and therapeutic efficacy in the brain.

We developed a triple combination formulation (TCF) that delivers the HDACi vorinostat (Vo) across the BBB to promote histone acetylation in the brain ^22^. Once-weekly administration over 3 months in a murine model of Niemann Pick Type C disease, increased brain *Npc1* transcript and protein levels, preserved neurites and Purkinje cells, reduced inflammation and delayed neurodegeneration and death ^22^. The TCF was well tolerated, with normal mice showing no metabolic toxicity, detriment in major brain neuronal or immune cells or *Hdac* gene-expression, even after 10 months of chronic, once-weekly exposure ^22, 23^. Vo is a broad spectrum HDACi^24^, and this raised issue about long term exposure and tolerance of TCF suggesting need to understand the specificity of repeated, long term TCF administration on target histones as well as other molecular and cellular responses in the brain.

Here we examined TCF-treatment in a mouse model of Kabuki Syndrome (KS), a monogenetic disorder, caused by heterozygous loss-of-function mutations in lysine-specific methyltransferase 2D (KMT2D) or lysine-specific demethylase 6A (KDM6A)^25, 26^. KMT2D trimethylates histone 3 lysine 4 (H3K4) to yield H3K4me3 ^27^, a mark of open chromatin. KDM6A demethylates trimethyl histone 3 lysine 27 (H3K27me3) ^28, 29^, a mark of closed chromatin. Both genes promote the opening of chromatin and gene expression, suggesting that drugs (like HDACi) that elevate open chromatin states may rescue deficit in hippocampal neurogenesis and function. Prior studies reported reduction in post-natal neurogenesis in the subgranular zone (SGZ) of the dentate gyrus (GS) in the hippocampus of a mouse model of KS that could be reversed by the HDACi AR-42 ^30^. AR-42 also showed effects on peripheral blood DNA methylation^31^. Defects of KMT2D in B cells ^32, 33^, cardiac, muscle, adipose and epithelial tissues ^34–36^ have been linked to immune disorders, heart and other systemic tissue diseases as well as cancers ^37^. However, the effects of reduction of KMT2D on brain immune cells and their contribution to hippocampal function are unknown.

Intellectual disability associated with functional brain deficit ^38^ presents as a major challenge in KS patients, for which treatments remain elusive. Intellectual disability arises from additional genetic diseases that control chromatin states ^39^ and can affect sizeable numbers of children world-wide ^40^. In this study, we combined a monogenetic mouse model of intellectual disability and response to HDACi with single-cell RNA sequencing (scRNA-seq). We reveal an original, unforseen defect in a major, brain immune cell type called microglia in KS and present evidence, its novel differentiation without activation that may treat neural disease in mice.

## Results

### Reduction of the open chromatin mark H3K4me3, hippocampal post-natal neurogenesis and microglia in the brain of *Kmt2d^+/βGeo^* (Kbk) mice

In the Kmt2d deficient mouse the catalytic SET (suvar, enhancer of zeste, trithorax) domain in one allele is replaced by a β Geo cassette (Kmt2d+/βGeo; Fig. 1Ai). Mice backcrossed to 99-100% to C57BL/6J background (see Materials and Methods) were confirmed for loss of the SET domain in ∼50% of *Kmt2d* transcripts in the brain (Fig. 1Aii, Fig. S1A) and reduction in tri-methylation of histone 3(H3K4me3) (Fig. 1Aiii). Bjornsson et al reported that Kbk mice showed decrease in post-natal neurogenesis (as measured by staining for doublecortin X (DCX, a marker for neural precursors and immature neurons), in the subgranular zone (SGZ) of the dentate gyrus (DG) of the hippocampus, that could be increased by administration of a pan HDACi AR-42^30^. As shown in Fig. 1B and Fig. S1B, we too find that Kbk mice show a reduction in staining of DCX compared to WT counterparts. We administered the TCF (DMSO+PEG+HPBCD+Vorinostat (Vo)) in once weekly injections: this regimen was known to be brain permeant and well tolerated in long term administration in mice^22, 23^. However, even after three months, neither the TCF nor Vo (the active HDACi ingredient in TCF) increased DCX or stimulated post-natal hippocampal neurogenesis in Kbk mice (Fig. 1B and S1B).

**Figure 1.**
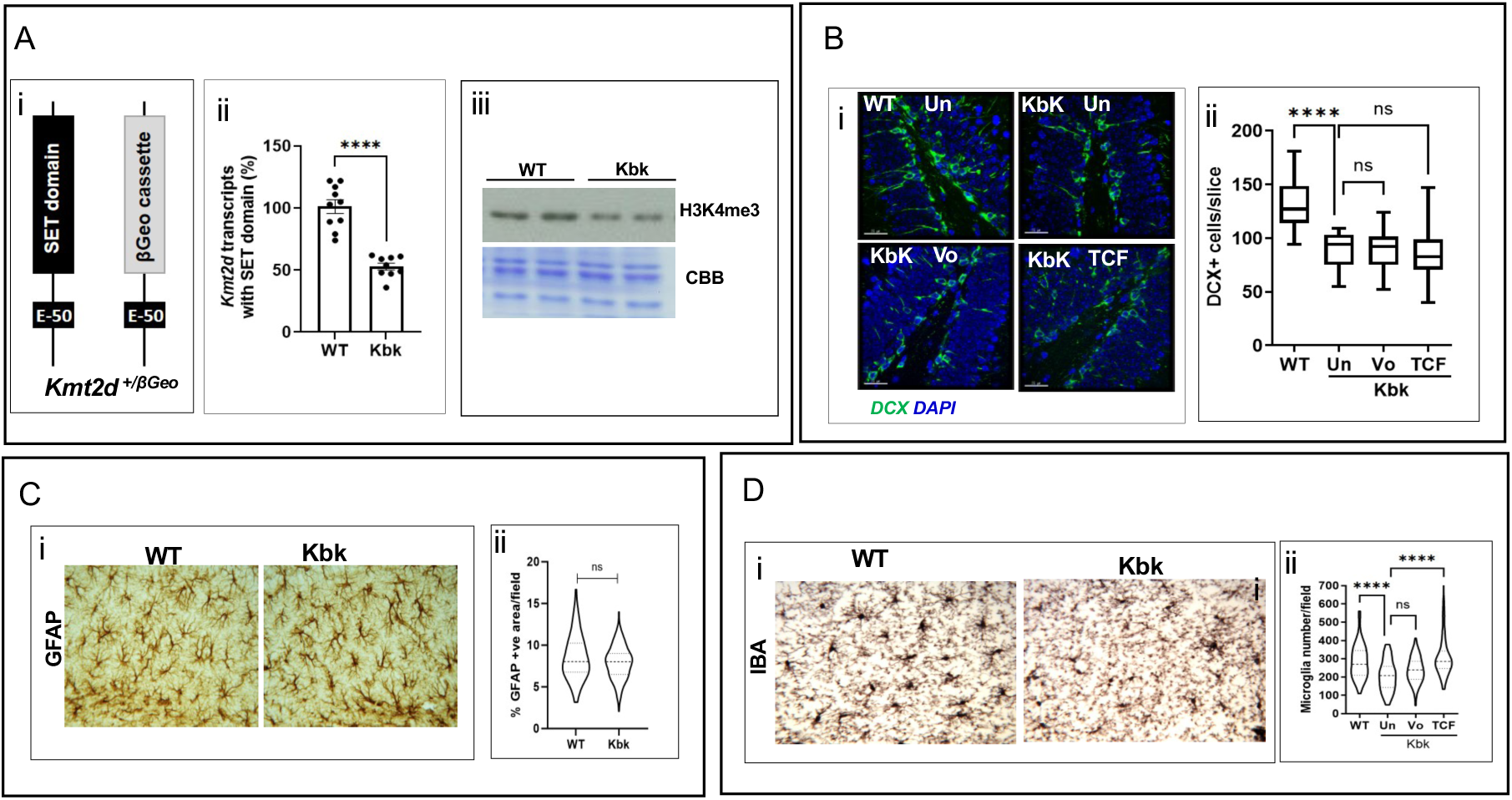
Effects of Kbk genotype on brain transcript levels, chromatin mark, post-natal hippocampal neurogenesis, astrocytes and microglia. **A. Comparison of transcript and the chromatin mark H3K4me3.** Schematic showing SET domain in one copy of the *Kmt2d* gene (coding region after exon 50) replaced with the βGeo cassette in the (Kbk) mouse model. **ii.** qPCR analysis of brain showing 50% of *Kmt2d* transcripts (measured by detection of exon 52) lacked the SET domain in Kbk relative to WT mice; n=9-10 (4-5F, 5M) per group**. iii**. Western blots of histone H3 lysine 4 trimethylation (H3K4me3) levels in the brain of Kbk mice. Equal sample loading shown by Coomassie brilliant blue (CBB). **B. Post-natal neurogenesis**. **i**. Micrographs show doublecortin (DCX, green) labeled neuroblasts in the SGZ of the hippocampal DG of WT and Kbk mice, as detected by indirect immunofluorescence assays. Mutants were un-injected (un) or injected with Vo (in DMSO+PEG) or TCF. Images shown are representative of 4-6 mice (2-3F, 2-3M, 2 months) per group. Blue indicates 4′,6-diamidino-2-phenylindole (DAPI) stain. Scale bar, 25 µm. **ii**. Quantification of DCX positive cells along the entire SGZ Brain sections (5 mm thick spaced at 40 mm) at 4-6 planes per mouse were analyzed; n=4-6 mice/group. Statistical analysis was conducted by one-way ANOVA with Tukey’s posthoc test. **C. Astrocytes**, **i**. Micrographs show GFAP positive astrocytes (brown) in hippocampi, as detected by immunohistochemistry. **ii**. Quantitation of hippocampal astrocytes after excluding granule cell regions and polymorph layer of the DG (to eliminate GFAP expressing neural progenitors). Percent GFAP positive area was calculated in 4-5 random fields at three depths (consistent across all mice) and the value from each image was used to plot the data. Statistical analysis by Mann-Whitney test. **D**. **Microglia. i.** Micrographs show IBA1 positive microglia (brown) in the hippocampus. **ii**. Quantitative analyses of hippocampal IBA1 positive microglia (under different conditions) are represented by Violin plots of images collected at 6-7 consistent planes (through the entire depth of the hippocampus). Statistical analysis was conducted by one-way ANOVA with Tukey’s posthoc test.

We next focused on the brain’s major immune cells, astrocytes and microglia, whose characteristics remain unknown in KS mice. As shown in Fig. 1C, astrocytes stained for their characteristic marker Glial Fibrillary Acid Protein (GFAP) showed no significant change in Kbk compared to WT mice (Fig. 1C; minor changes seen with individual genders in Fig. S1C, did not prevail when both were combined). We subsequently examined microglia, which are known to be present in higher density in the hippocampus^41^. As shown in Fig. 1D and Fig, S1D microglia were stained with their characteristic marker, IBA1 (Ionized calcium-binding adaptor protein-1 an actin-binding protein). Notably, in Kbk mice, the maximal and median microglial densities were lowered by 33% and 23% compared to wild type counternparts. TCF (but not its independent components) quantitatively increased microglia densities in the hippocampus (Fig. 1D, Fig. S1D). Together these data suggested that hippocampal microglia are reduced in Kbk mice. Moreover, although the TCF failed to stimulate post-natal neurogenesis in the SGZ of the DG, it may influence microglial function in the hippocampus.

### Single cell RNA sequencing of wild type and mutant microglia and their responses to chronic TCF-administration

To understand molecular changes in hippocampal microglia elicited by TCF, we compared single cell RNA sequencing of cells isolated from WT and Kbk mice in presence or absence of treatment. As summarized in Fig. 2A and Fig. S2A, a total of 60 mice were utilized (balanced for sex and genotype). This included a cohort of six mutant and six WT mice at one month of age that provided a baseline to compare early post-natal differences, as well as treatment groups of 24 WT and 24 Kbk at three months of age, that received weekly TCF or control treatments of PEG (in DMSO) and/or HPBCD (in DMSO+PEG). Single-cell preparations of isolated hippocampi were hash tagged (to distinguish between WT and Kbk; Fig. S2B-D) before advancing to single call capture, library preparation, RNA seq cell ranger and Seurat Analysis (Materials and Methods). After filtering, we obtained a total of 25,165 high-quality healthy single cells. These included 12,216 microglia (Supplementary Table S1), confirming their enrichment in our preparations. Uniform Manifold Approximation and Projection (UMAP) analysis was used to reduce the multidimensional scRNA-seq data into 28 distinct clusters that were categorized into 11 major cell types which included of microglia, endothelial cells, astrocytes, choroid plexus epithelial cells, oligodendrocytes, oligodendrocyte progenitor cells, dendritic cells, pericytes, neural progenitor cells/neurons, vascular leptomeningeal cells, and immune cells (Materials and Methods; Supplementary Table S1 and Fig. 2C). Microglia with 12,216 cells comprised 48.5% of total and were judged to be suitably abundant for further single cell analyses (while the remainder groups were not; Supplementary Table S1-S4). We pre-processed the microglial subsets before performing UMAP analysis, which revealed presence of seven clusters delineated computationally (0-MG to 6-MG) (Fig. 2D) suggesting they may span multiple states, across genotypes and treatments.

**Figure 2.**
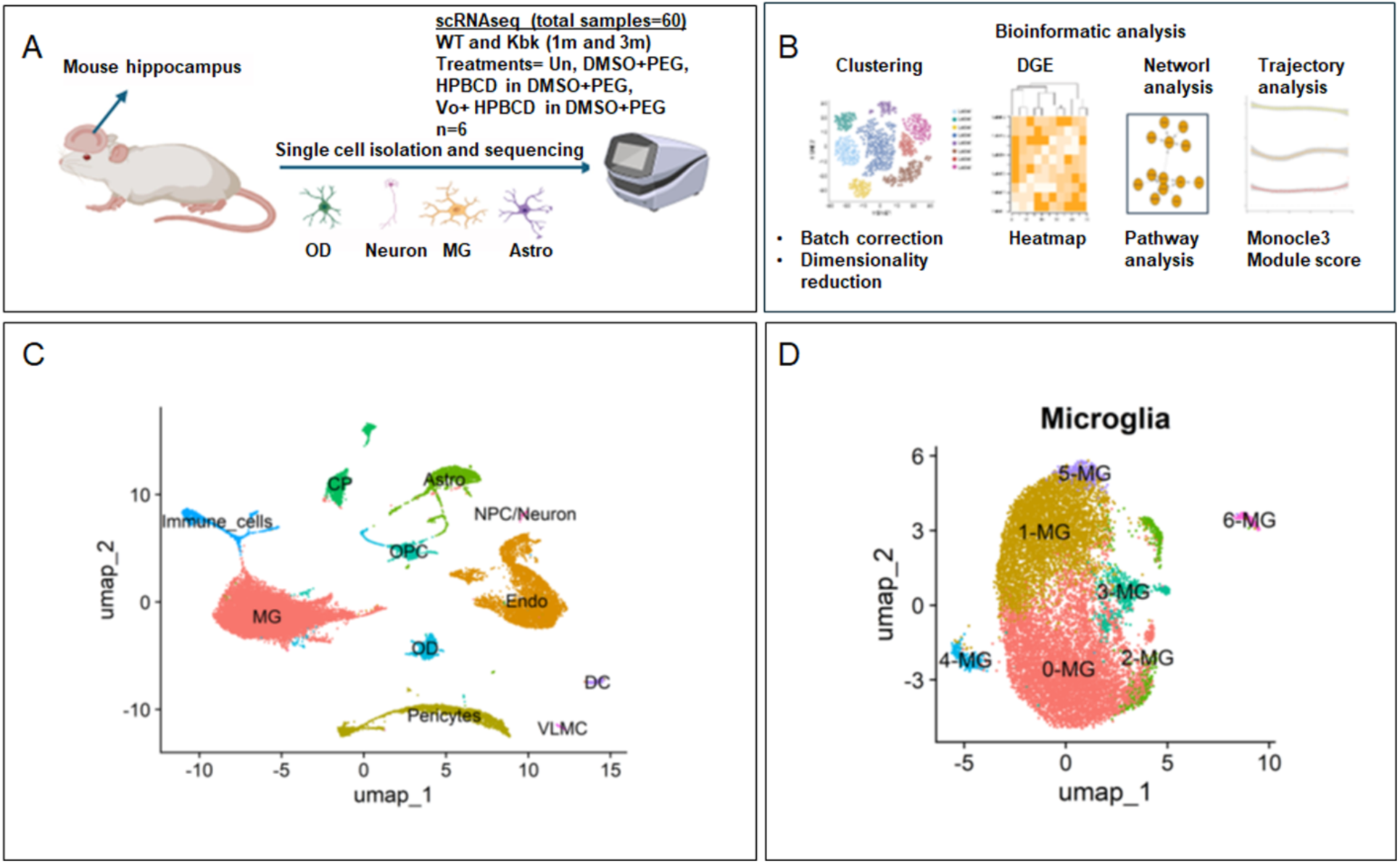
Single cell RNAseq analyses of gene expression changes seen in microglia isolated from mice hippocampus of WT and Kbk_ mice. **A** Graphic representation of overall design of single cell RNAseq experiments generated using BioRender. Briefly a total of 60 mice (WT and Kbk) were used. **B**. Details of samples, sequencing methods and downstream bioinformatic analyses. **C**. UMAP of 25165 cells which are clustered into ten different sub-cell types; **D**, The UMAP represents 12,216 microglia cells that could be computationally clustered into seven different microglial subsets (0-MG to 6-MG).

### Downregulation of hippocampal microglia seen in Kbk mice is mitigated by TCF

We initiated differential gene expression (DGE) analysis to investigate changes in transcriptional signatures of microglia across experimental groups. Due to the sparsity and high noise inherent in scRNA-seq data, we did not use traditional DGE approaches, but rather a pseudobulk analysis, which aggregates single-cell expression counts from individual samples, effectively transforming the data into a bulk RNA-seq format.

Comparisons of DGEs of untreated WT and Kbk mice at three months are shown in Fig. 3A. Here three female and one male Kbk sample formed a distinct cluster, while the remaining two males grouped more with WT counterparts. This type of heterogeneity is characteristic of defect in Kmt2d, since the extent of loss of chromatin and consequences for gene expression, can vary between mutant animals. The major cellular pathways impacted were investigated using Ingenuity Pathway Analyses (IPA; Fig. 3B; Fig. S3B-S3C). Of the top six predicted pathways, five suggested global reduction in translation and oxidative phosphorylation (Fig 3B). Notably Aif 1(the gene coding for the major microglial marker IBA1) was in the top ten downregulated transcripts identified (p < 0.05, fdr <0.05; Fig. 3Ci). IBA1 was also confirmed to be reduced in hippocampal Kbk microglia detected by immunohistochemical (IHC) which were additionally reduced in soma size (Fig. 3Cii-iii). Thus, at 3 months, Kbk microglia appear to be arrested forms, small and downregulated in basic cellular pathways and suppressed in a major proliferative, inflammatory maker IBA1. Examination of one month old mice also showed heterogeneity in microglia, but at this age the major changes in mutants were in signaling and vesicular pathways (Fig. 3D-E). Together, the data in Fig. 3A-E suggest that changes in hippocampal mutant microglia emerged as early as one month after birth but global reduction in cellular translation, energy production and IBA1 and progression to small, arrested forms. This is in contrast to findings that in normal mice early microglia display a round amoeboid morphology, active proliferation and phagocytosis^42^

**Figure. 3.**
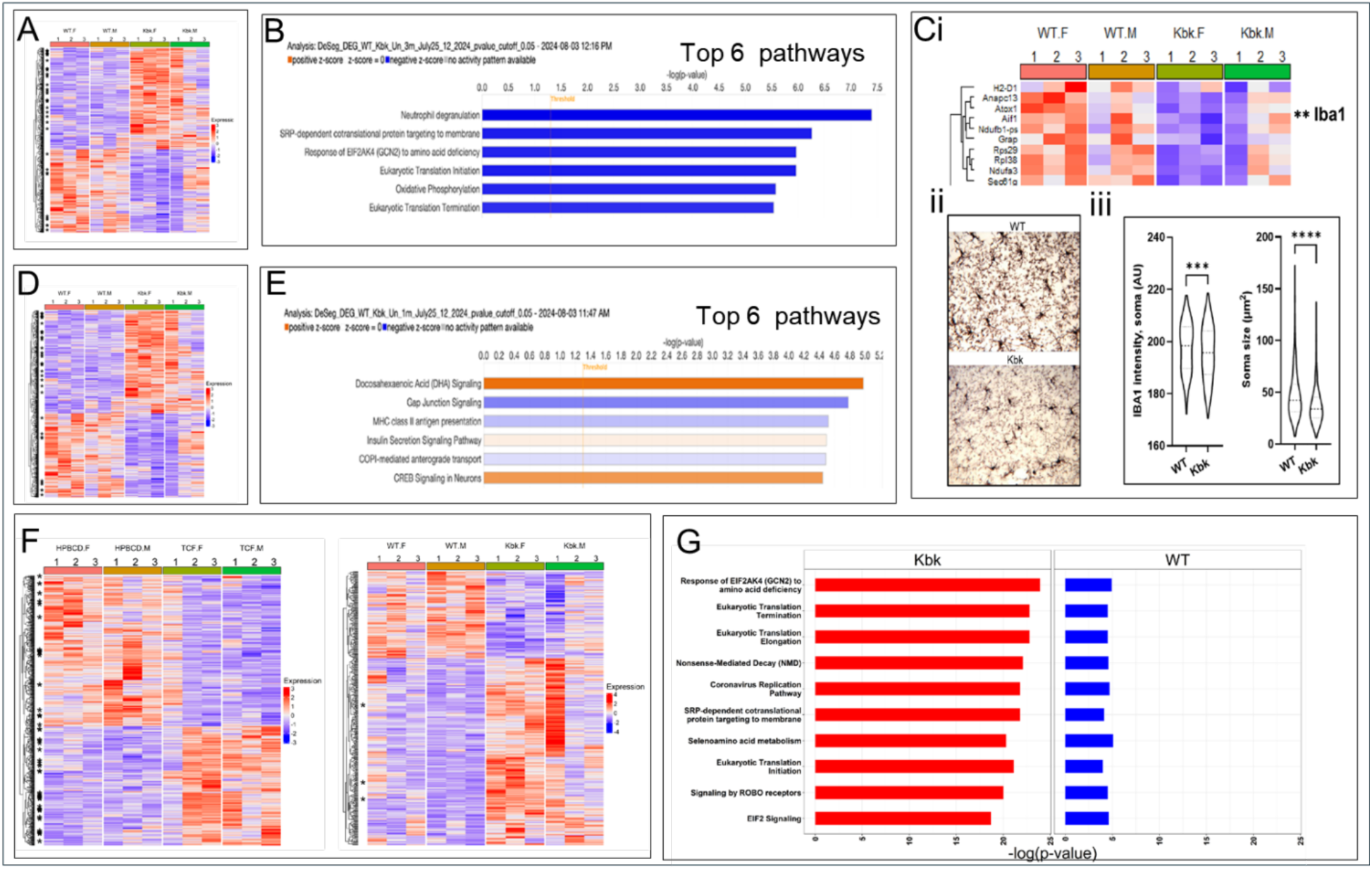
Comparative change in microglial gene expression, predicted pathways and cellular phenotypes in Kbk and WT, dependent on age and/or TCF treatment. **A-C.** 3-month untreated mice. **A.** Differential expression of genes (DEG) in Kbk versus WT mice, with p-value < 0.05; asterisks indicate fdr < 0.05. **B.** Bar plot of top 6 pathways predicted to be down regulated (blue) in Kbk by IPA. **Ci,** DEG of top 10 genes from A (red upregulated in Kbk versus WT); **C ii-iii,** IHC of Iba1 stained microglia and associated quantification of soma intensity and size by Imaris. **D-E.** 1-month untreated mice. **D**. DEG with p-value < 0.05: asterisks indicate fdr < 0.05. **E.** Top 10 pathways predicted to be downregulated in Kbk vs WT by IPA. **F-G.** Effects of TCF-treatment versus vehicle control (HPBCD) administered once weekly for three months. **F.** DEGs with p-value < 0.05: asterisks indicate fdr < 0.05. **G** Bar plot of top 10 pathways predicted by IPA based on DEG responses in **F**. In panels **A-Ci, D-G**, red and blue respectively show relative up and downregulation of genes or pathways.

To examine the effects of TCF, mice were first administered at one month of age and followed up weekly for up to 3 months. As shown in Fig. 3F, by DEG analyses, TCF treatment improved clustering in five out of six Kbk mice relative to control HPBCD-treated samples. We also observed a clustering pattern emerging between DMSO+PEG and HPBCD (in DMSO+PEG) treatments (Fig. S3A). But these were distinct from TCF-induced clustering seen in treated mutants. IPA analyses (Fig. 3G) suggested that in Kbk, TCF treatment globally stimulated translation and other cellular pathways, whereas in WT mice, TCF mildly suppressed a small subset of genes and pathways (3F-G). Together these data suggest that in Kbk, TCF effectively reversed massive transcriptional down-regulation associated with three-month old mutants but WT with normal heterochromatin were protected from TCF.

### Chronic TCF administration selectively stimulates microglial histone H3 and overall increase in H3K4 open chromatin mark in Kbk brain

Histone proteins in the nucleus are known to play a key role in the maintenance of chromatin. Vorinostat (Vo), the HDACi in TCF blocks deacetylation of histones by targeting multiple HDACs^43^. IPA analyses predicted elevation of the chromatin mark H3K4me3 in Kbk, without stimulation of Kmt2d but unexpectedly driven by increase of a single histone H3 (Fig. 4A; Supplementary Fig. S4). Importantly, the other histones (1, 2 and 4) and associated pathways were effectively unperturbed, suggesting chronic TCF resulted in selective, long-term targeting through histone H3.

**Figure 4.**
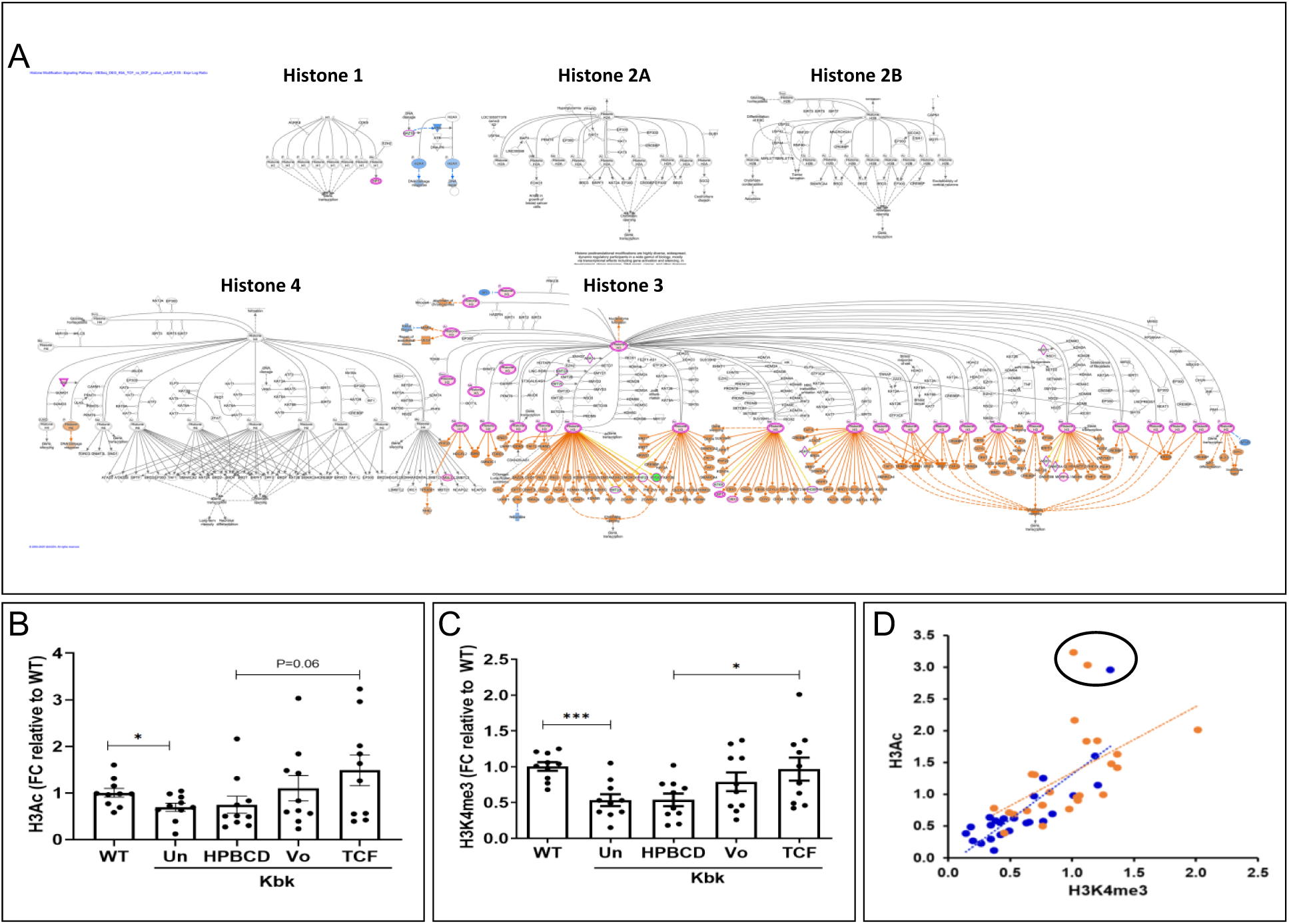
TCF targets Histone H3 dependent increase of open chromatin independent of *Kmt2d* gene as predicted by IPA and validated in mouse brains. **A**. IPA analyses of DEG shown in Fig. 3G, suggest that in Kbk mice, weekly TCF administered over 3 moths, targets a single histone H3 (red; p < 0.05), acetylation (H3Ac) of which is predicted to stimulate opening of chromatin (orange) independent of Kmt2D (black box). Elevation of H3 may reflect increased life span of the S phase the time of greatest histone transcription^64^. **B-C.** Validation of TCF-mediated H3Ac and H3K4me3 in whole brain. Quantitative analysis of western blots showing fold change (FC) in (**B**) H3Ac or (**C**) H3K4me3. Kbk mice were given or its components as indicated, starting at P21-P23, and the brain was analyzed at ∼3 months of age. All data were normalized for total histone protein and fold change is relative to the average of wild type (WT) mice. n=10 (5F, 5M)/experimental group. **D**. Correlative analyses of levels of H3Ac and H3K4me3 in brains of 50 mice. Pearson correlation co-efficient (r)=0.7, *P< 0.0001*. Data is separated by gender (males, blue; female red). Pearson correlation co-efficient (r) males, r=0.8*, P<0.0001*; females, r=0.5, *P=0.008*., Un, untreated. *P* values by unpaired, two-tailed student’s t-test. *\*P<0.05,**P<0.001, ****P<0.0001*

As further validation, we examined the utility of TCF to stimulate histone H3 acetylation and the open chromatin mark in the whole brain. As shown in Fig. 4B, after 3 months of weekly injections, TCF appeared to increase median values of H3Ac levels in the Kbk brain compared to HPBCD (in PEG+DMSO; that lack HDACi (Vo; but with a p-value of 0.06). Vo (in DMSO+PEG) may also have an effect but based on H3Ac levels alone, we were unable to draw a firm conclusion on the significance of TCF-induced acetylation. No sex specific effects were seen (Supplementary Fig. S5) However, as shown in Fig. 4B, TCF induced a significant increase of the open chromatin mark H3K4me3, relative to HPBCD (in PEG+DMSO). Vo (in PEG+DMSO) also increased levels of H3K4me3 but with intermediate values that could not be separated as statistically significant from TCF or HPBCD (in PEG+DMSO). Notably (and as shown in Fig 4C), in >90% of animals there was a strong positive correlation between acetylation and trimethylation which was seen in both genders (Fig. 4D). Only three of the fifty mice showed disproportionately elevated levels of acetylation. Together, these data established a primary effect of histone H3 mediated acetylation-dependent promotion of H3K4me3 in the Kbk brain. TCF achieved higher levels of acetylation and trimethylation than its components, suggesting its brain permeant and HDACi-dependent properties were important for acetylation-dependent histone trimethylation throughout the brain.

### TCF Treatment Drives Divergent Microglial Trajectories in WT and Kbk mice

The distinct morphological differences observed in microglia from Kbk and WT mice (in Fig. 3) led us to hypothesize that these variations reflect underlying molecular differences in cell states. To investigate this, we revisited the scRNA-seq dataset and applied pseudotime analysis, a computational technique that organizes cells along a theoretical trajectory, representing their progression through biological processes. Pseudotime analysis enables the reconstruction of temporal events at a single-cell resolution, providing a deeper understanding of complex cellular dynamics^44^. Using the R package Monocle3, we constructed a microglia trajectory from 12,216 cells across all experimental groups, assigning pseudotime values to each cell. Cells at the beginning of the trajectory represent earlier, less differentiated states, while those at later stages represent more advanced differentiated states. To quantify these trends, pseudotime values were divided into 25 bins, and the proportion of cells from each experimental group was calculated within these bins^45^. WT untreated-mice at 1 month of age were taken to be the start of the trajectory.

As shown in Fig. 5A-B, microglial trajectories exhibited distinct patterns between Kbk and WT groups under different treatment conditions. For Kbk mice, the proportion of cells treated with PEG (+DMSO), HPBCD (in DMSO+PEG), and TCF decreased along the pseudotime trajectory, with Pearson correlation R = −0.12, p = 0.57; R = −0.44, p = 0.03; R = −0.58, p = 0.0028, respectively). In contrast, WT mice treated with PEG (+DMSO), HPBCD (in DMSO+PEG), and TCF showed increased proportions along the pseudotime trajectory (Pearson correlation R = 0.71, p = 0.00087; R = 0.68, p = 0.0014; R = 0.76, p = 4.7e−05, respectively). These opposing trends highlight a fundamental divergence in how WT and Kbk microglia progress through their cellular states.

**Figure 5.**
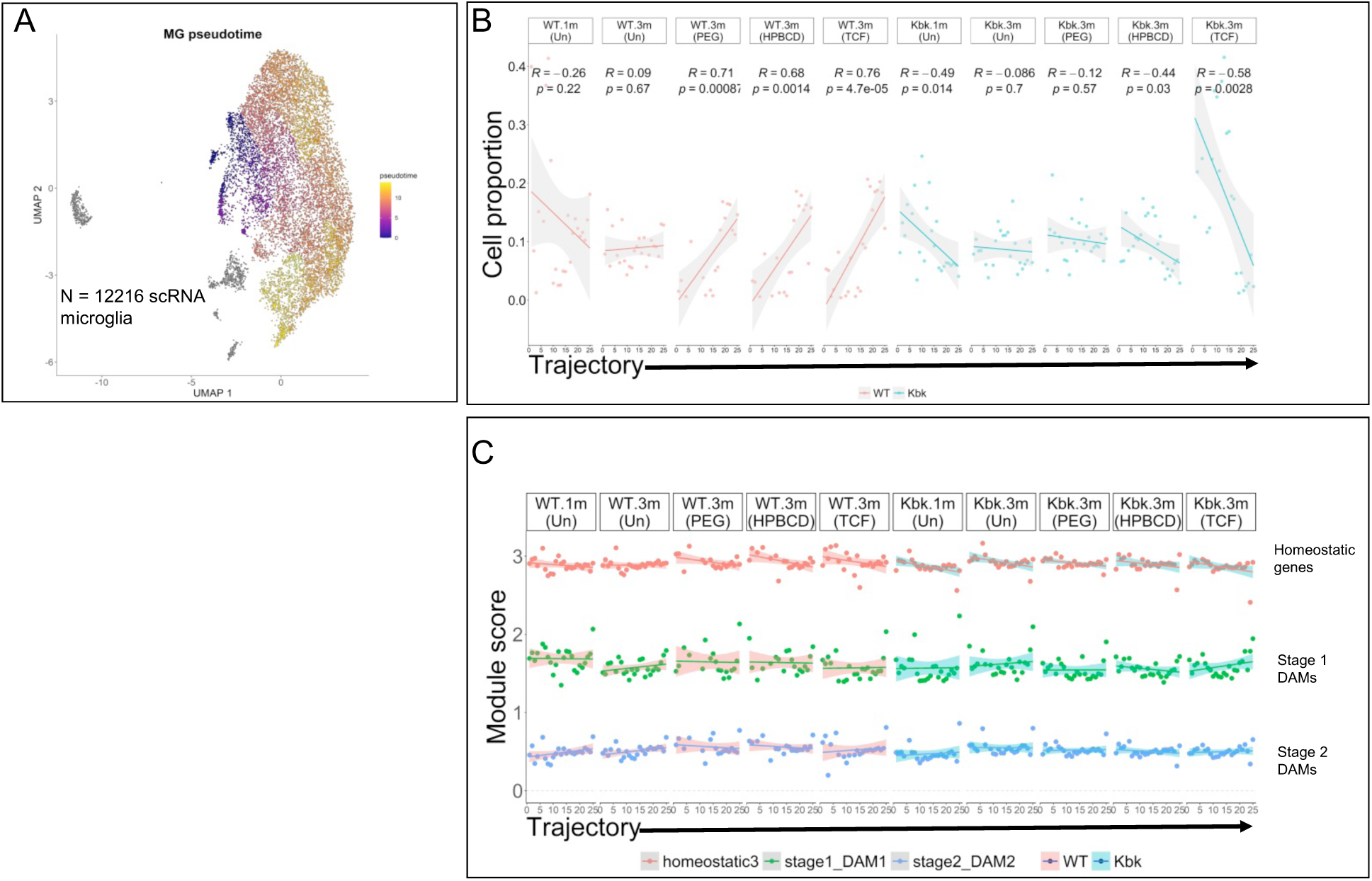
Cell trajectory analyses using pseudotime analyses tracking homeostatic and disease markers. **A**. This UMAP plot shows the dimensionality reduction of microglia derived from a scRNA-seq analysis of 12,216 cells. Each cell is represented as a point in the plot, with colors corresponding to their pseudotime values. The smooth color gradient across the UMAP indicates a progression of cellular states. **B**.This scatter plot illustrates the distribution of microglia cells across 25 equal intervals (bins) along the trajectory. The solid line represents the linear regression fit, indicating the relationship between cell proportion and trajectory. The shaded gray area surrounding the line represents the 95% confidence interval (CI), giving a visual of the statistical certainty of the regression line. The Pearson correlation coefficient (R) and p-value from a two-sided statistical test are displayed, indicating the strength and significance of the correlation between the variables. **C.** This scatter plot illustrates the correlation between module scores and pseudotime (trajectory) for different gene modules, specifically the Homeostatic (Hexb, Tmem119, Cx3cr1, P2ry12, P2ry13, Cst3, Cd33, Csf1r, Ctss, Tmsb4x, and C1qb), Stage 1 DAM (Tyrobp, Ctsb, Apoe, B2m, Fth1, Trem2, and Ctsd) and Stage 2 DAM (Lpl, Cst7, Axl, Itgax, Spp1, Cd9, Ccl6, Csf1, Trem2, Timp2 and Ctsl) gene modules. The solid lines show less regression for each gene signature, and the shaded gray areas around them represent the 95% confidence intervals (CIs). These regressions help visualize how the module scores change across the cell trajectory.

While different treatments did not significantly alter the linear relationship between pseudotime and cell proportions in WT microglia, TCF exerted a marked effect on Kbk microglia (Fig. 5B). Specifically, TCF led to a significant decrease in cell proportions along the pseudotime trajectory compared to HPBCD treatment. This effect suggests that TCF influences the progression of Kbk microglia to drive their differentiation toward earlier, distinct cellular states and outcomes (driving toward the start of the trajectory set at WT at one month). In contrast, WT microglia initially appeared in lower proportions at early pseudotime stages, but their numbers increased as pseudotime progressed, indicating a gradual advancement (without differentiation) toward later states.

### Module score vs pseudotime analysis suggested that the distinct molecular gene modules responsible for microglial activation were unchanged by TCF or its components

In a wide range of neurodegenerative disorders such as Alzheimer’s disease, amyotrophic lateral sclerosis, and multiple sclerosis, the expression of both stage 1 and stage 2 disease associated microglia (DAM) genes, as well as homeostatic genes, is differentially regulated ^46^. DAM arise from homeostatic microglia, which normally play a crucial role in maintaining the health of the central nervous system (CNS) by clearing debris and supporting neural circuits. The shift from homeostatic microglia to an activated DAM state is a key process in neuroinflammation by upregulation of genes associated with inflammation, lipid metabolism, and phagocytosis (such as Lpl, Cst7, and Axl). The DAM is associated with increased expression of DAM gene modules (Stage 1 DAM genes: - Tyrobp, Ctsb, Apoe, B2m, Fth1, Trem2, and Ctsd, Stage 2 DAM genes:- Lpl, Cst7, Axl, Itgax, Spp1, Cd9, Ccl6, Csf1, Trem2, Timp2 and Ctsl) and decreased expression of homeostatic genes (Hexb, Tmem119, Cx3cr1, P2ry12, P2ry13, Cst3, Cd33, Csf1r, Ctss, Tmsb4x, and C1qb)^47, 48^. The DAM cells are activated by Trem2 signaling pathway and this activation leads to the transition from homeostasis to neuroinflammation. Fig. 6C illustrates the module score trends along the pseudotime axis across the different experimental treatments in Kbk and WT mice. From our module score vs pseudotime analysis we can say that there is no major difference in homeostatic, stage 1 and stage 2 DAM gene modules across any treatments, therefore, implying that at the molecular level, relative to wild type, microglia activation or microglial inflammation is not aggravated in the hippocampus of Kbk mice in their untreated state or in different treatments, including the TCF that has the greatest effect in influencing the progression and cellular state of Kbk microglia (Fig. 5A-B)

**Figure 6:**
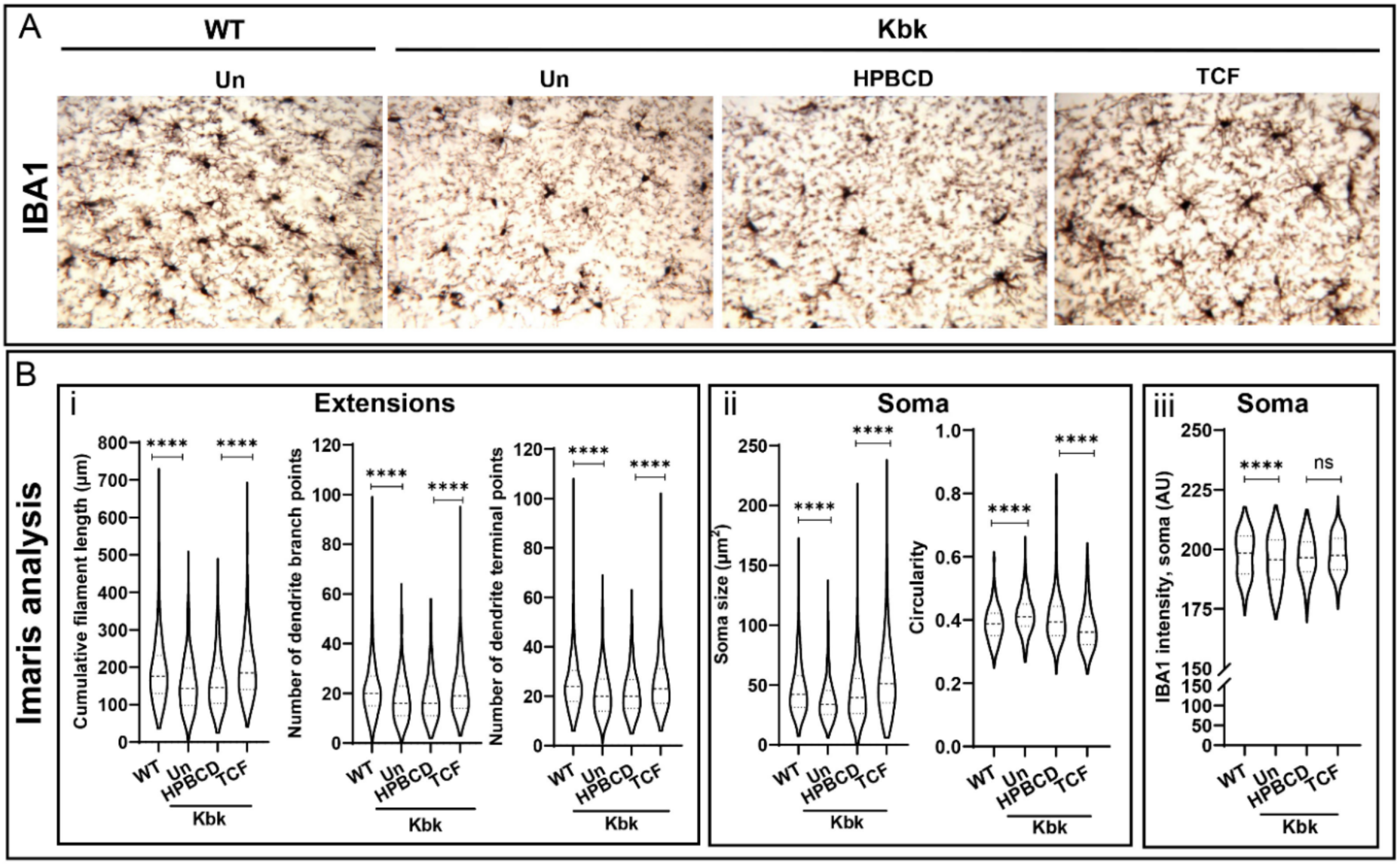
Defects in microglia in the hippocampus of KS mice and their response to TCF. **A.** Micrographs show density of IBA1 positive microglia (black) in the hippocampus. Treatments are as indicated. Representative images collected at 6-7 consistent depths (through the entire depth of the hippocampus) with 40x objective are shown. **B.** Violin plots of Imaris-based quantitative data of microglial filament length, number of branch and terminal points assessed from 400-450 hippocampal microglia (40±10 per mouse) per group. (B, middle panel) microglia soma size and circularity and (B, right panel) mean IBA1 intensity, as assessed from 600-700 (60±15 per mouse) microglia per group. For B, microglia were taken from two random fields at two consistent depths of the hippocampus in each mouse. Statistical analysis in B panels was undertaken by nonparametric Kruskal-Wallis test. In all panels we used n=8-9 mice (4-5 females, 4 males) per group at 3 months of age. WT wild type, Kbk Kabuki, Un Untreated, HPBCD Hydroxyl-propyl-cyclodextrin, TCF triple combination formulation. **P<0.01, ***P<0.001, ****P<0.0001.

### Impact of TCF on microglial density, filamentous extensions, soma size and overall surveillance phenotypes in the hippocampus of Kbk mice

Microglia are resident macrophages of myeloid origin in the brain, characterized by filaments/extensions that are connected to a cell body (or soma). Lengthening and branching of their filaments dominate at homeostasis to enable sensing and surveillance of brain areas^49^. Activation of microglia in response to infection or injury^50^, results in shortening of extensions and enlargement of soma, to promote phagocytic functions and reduce surveillance. Ameboid morphologies are associated with neuroinflammation and ageing^51^.

Since morphometric characterization is facilitated by automation, we utilized Imaris software to undertake quantitative, high-throughput analyses of 4-500 IBA1-stained hippocampal microglia from 3-month-old WT, Kbk animals and Kbk animals treated with TCF and appropriate control treatments. HPBCD (in DMSO+PEG), Vo (in DMSO+PEG). Compared to wild type (Fig. 6A) in Kbk mice (Fig. 6Bi), the maximal filament extension length, numbers of dendrite branch points and terminal points were reduced by 30-36%, while median levels of these parameters showed reduction of 17-20%. Maximal and median soma sizes in Kbk were also decreased (by 20%) while circularity showed a smaller increase by 6-8% (Fig. 6Bii). TCF promoted length and numbers of filament branch and terminal points by 42-64%. It increased the maximal and median size by 10-30% and reduced maximum and median roundness by 25 to 8%, respectively. In contrast, HPBCD (in PEG+DMSO) or Vo (in PEG+DMSO) had minimal or no significant effect (Fig. S7A and S7B), suggesting that only TCF specifically induced major changes in filament parameters and influenced soma (Fig. S7). The differences in IBA1 levels in mutants compared to wild type was 1-2% across groups, suggesting neither mutation nor treatments led to activation (Fig. 6B right panel and Fig. S7A upper right panels and S7B lower right panels). These changes were seen in both genders (Fig. S7) but with slight differences. The effects on soma size and circularity were variable between genders. In males, HPBCD (in PEG+DMSO), Vo (in PEG+DMSO) and TCF increased soma size to the untreated WT level, but the magnitude of TCF effect was maximal. In females, however, only TCF caused an increase in soma size (Fig. S7A and S7B) and soma size was reverted to WT levels by TCF. The opposite trend was observed for circularity (Fig. S7A and S7B, lower middle panels).

To better visualize the above changes, rendered Imaris models were used to display the range of extensions of hippocampal microglia from wild type mice and mutant mice, as well as the effects of different treatments (HPBCD (in DMSO+PEG), Vo in DMSO+PEG) and TCF) (Fig. 7). Wild type mice showed microglia with the highest level of ramified extensions branch and terminal points (High). Wild type mice also displayed median and low levels of filaments. The greatest filament length, branches, and end points of Kbk mice were comparable to the wild type ‘median’ group. Of the treatment groups, TCF-induced substantial increase in filament length, branch, and endpoints. These effects were seen in both genders and suggested that in conjunction with decrease in density, there may be reduced surveillance functions of hippocampal microglia in Kbk mice, and both can be restored by TCF (Fig. 6 and 7).

**Figure. 7.**
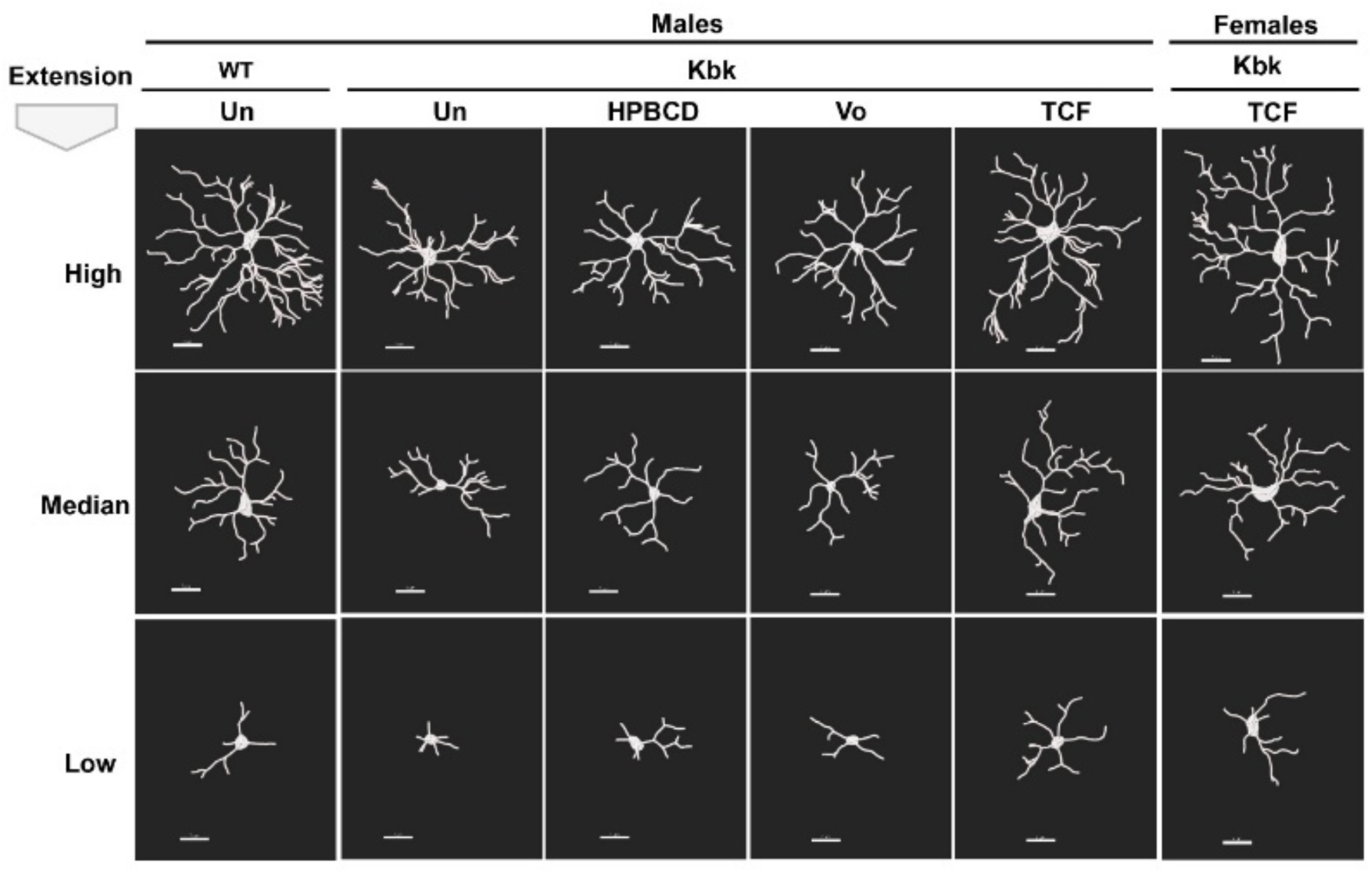
Imaris renditions of IBA stained microglia: effect TCF on surveillance phenotypes in hippocampus of Kbk mice. Rendered Imaris models showing extensions and soma in hippocampal microglia in wild type mice and *Kmt2d+/βGeo* untreated or treated with TCF or its components (as indicated). Representative models of microglia (based on 200-300 microglia per group, ∼40-75 per mouse) from males, show characteristics of branched and long extensions (90-100 percentile, high), intermediate size branches and ramification (median) and small filaments with fewer branches (1-10 percentile; low). Females show the same trends (not shown) but representative models of microglia from TCF-treated females are shown for comparison.IBA1-stained images of brain sections were taken from two random fields at two different depths in each mouse (n=8-9, 4-5 females, 4 males; ages 3 months). WT wild type, Kbk Kabuki, Un Untreated, HPBCD Hydroxyl-propyl-cyclodextrin, TCF triple combination formulation. Scale bar, 7µm.

### Effects of administration of TCF on long term hippocampal functions in Kbk mice

The Novel Object Recognition (NOR) test is a well-established and widely used measure of learning and memory in mice^52^ (and in prior work has been used to assess Kbk mice). Kbk animals showed reliable memory and learning defects as measured by a significant reduction in the Discrimination Index (DI) compared to wild type mice in the NOR test (Fig. 8A). DMSO, DMSO+PEG and Vo (in DMSO+PEG) had no significant effect. TCF significantly increased the DI to levels seen in wild type mice (Fig. 8A). Increased DI was also seen in both genders of TCF-treated mice (Fig. S8A).

**Figure 8:**
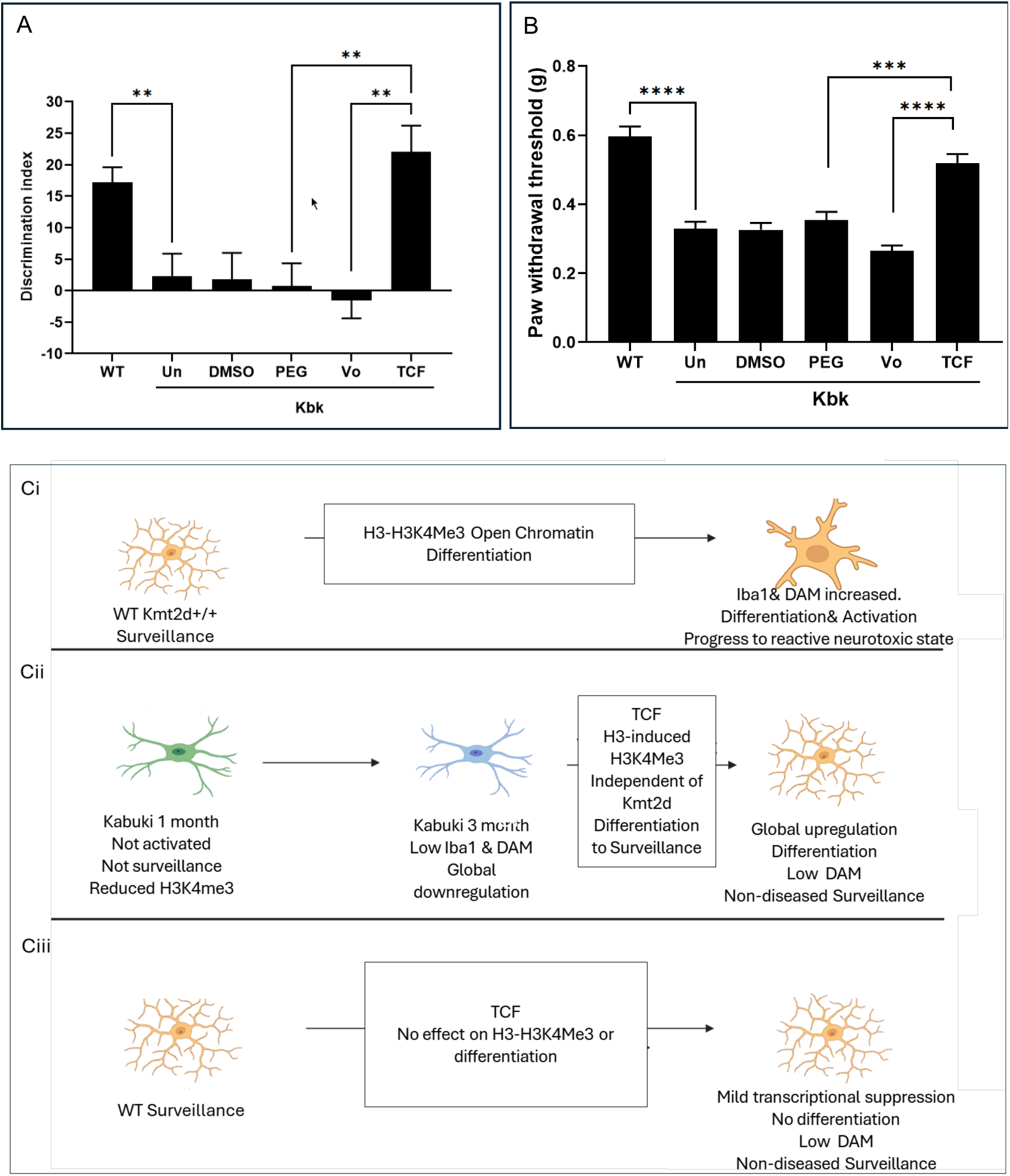
TCF-induced behavioral responses and a model H3K4m3-microglial differentiation without activation in the Kabuki hippocampus. **A**. Discrimination index (DI) was determined using the NOR assay to assess the hippocampal function of learning and memory of Kbk mice treated with TCF or its components as indicated. Mice were assessed at 5-7 months of age. Numbers of mice in different groups are WT, n=25 (12F, 13M), Kbk left un-injected (un, n=24, 10F, 14M) or injected with DMSO (n=10, 5F, 5M), PEG with DMSO (n=12, 6F, 6M), HPBCD in PEG and DMSO (n=20, 11F, 9M) or TCF (n=18, 9F, 9M). Data are mean ± SEM, one-way analysis of variance (ANOVA) with Tukey’s posthoc test. **B** Bar diagram showing paw withdrawal threshold, an indicator of tactile sensitivity assessed by the Von Frey (VF) assay. Both the left and right hind paws of each mouse were assessed on three consecutive days and average values were used for plotting. Numbers of mice and ages were, WT, n=41, 19F, 22M, 3-7 months), Kbk left un-injected (un, n= 38, 17F, 21M, 3-7 months) or injected with DMSO (n=26, 10F, 16M, 3-7 months), PEG with DMSO (n= 24, 12F, 12M, 3-7 months), Vo in PEG and DMSO (n=14, 7F, 7M, 3-4 months), HPBCD in PEG and DMSO (n=47, 21F, 26M, 3-7 months), or TCF (n=51, 25F, 26M, 3-7 months). Data are mean ± SEM, nonparametric Kruskal-Wallis test. **C. Model for TCF-induced H3K4m3-microglial differentiation without activation in Kbk mice**. **Ci.** In most neurodegenerative disorders, homeostatic microglia in response to increasing the open chromatin mark H3K4me3, differentiate and become activated to express elevated levels of Iba1 and other disease-associated microglia markers (DAMs). Activated microglia further transform into reactive microglia that display high levels inflammatory secretions and neurotoxicity. **Cii.** Kbk microglia, which are neither in activated nor surveillance state, undergo post-natal reduction of inflammatory markers and near global shut down of transcription from 1-3 months. TCF and H3-mediated increase of H3K4me3 open chromatin mark induced microglial differentiation, but not activation, rather differentiated microglia persist in a non-diseased surveillance state which confers behavioral benefit in mice. **Ciii**. In response to TCF, wild type homeostatic microglia do not undergo differentiation, but remain as non-diseased surveillance types with mild transcriptional suppression, suggesting wild type, normal heterochromatin protects against TCF-induced activation and differentiation.

For a second behavioral assay, we explored tactile sensitivity/nociception since work suggests this is also a hippocampal function^53–55^ and tactile avoidance is reported in KS patients^56, 57^. We therefore assessed mice for tactile responses using von Frey filaments^58^. As shown (Fig. 8B), wild type mice displayed 50% paw withdrawal threshold of 0.61g, which was reduced to 0.33 g in Kbk. DMSO, PEG (+DMSO) had no effect compared to controls. Vo (in PEG+DMSO) also had no effect, but TCF treatment improved the paw withdrawal threshold to 0.5g. These effects were seen in both genders (Fig. S8A and S8B) establishing tactile hypersensitivity as a measurable symptomatic domain in KS mice. Moreover, TCF (but not its components) improved tactile responses to within 80% of that shown by wild type mice. It was not due to increased levels of *Kmt2d* transcript or the H3K4me3 mark (data not shown), suggesting independence of Kmt2D.

## Discussion

We provide the first evidence for defects and disease in microglia in Kabuki syndrome in mice. Microglia are known to become diseased in Alzheimer’s, and other poly- and monogenetic neurological disorders as well as in cerebral infections. These microglia are highly activated, display disease associated microglia markers (DAMs) cause inflammatory pathology and become neurotoxic in the brain^59–62^. Histone H3-dependent stimulation of the open chromatin marker H3K4m3^63^ is well known to promote activation of microglia and their progression to diseased, neurotoxic states (see model in **Fig. 8Ci**). Surprisingly, diseased Kbk microglia are repressed in basic cellular functions of protein production and respiration, do not present capacity for inflammation, and even at three months post-natal, fail to display DAMs (summarized in **Fig. 8Cii**). TCF treatment drives diseased Kbk microglial trajectories towards differentiated states enriched for the H3K4me3 mark, suggesting that the TCF acts by driving open chromatin states to propagate differentiation of small-arrested Kbk microglia into surveillance forms, but in absence of increasing DAMs (**Fig. 8Cii**). Hence we propose prevention of DAM increase after elevating H3K4me3 shown in Fig. 8Cii, may be a pathway of generating non-diseased surveillance forms in the brain.

Our studies also underscore the specificity and therapeutic relevance of histone H3 modifications in addressing chromatin dysregulation. Although Vo can inhibit multiple HDACs, our finding that TCF selectively increases levels of histone H3, suggests that in microglia, it may preferentially inhibit HDAC3. Strong stimulation of the H3K4me3 mark upon HDAC3 inhibition has been shown to be associated with microglial differentiation and proliferation, but as indicated earlier in most neurodegenerative diseases this is associated with inflammation (**Fig. 8Ci**) ^63^. In our studies in Kbk mice weekly administration of the TCF enables short exposure (2-4h) of HDACi (2-4 h) with intermittent seven-day rest periods which may be important for selective action through histone H3. Although acetylation is not necessarily expected to increase levels of the target histone per se, small increases in the length of the S phase^64^ during microglial differentiation and proliferation may increase levels of histone H3. This may be amplified after repeated exposure to TCF and may further contribute to TCF-induced increase in H3K4me3. TCF’s induction of a large transcriptional response with positive amplitude in Kbk but not WT microglia suggests that open chromatin, may shield against aspects of inhibition of HDAC3 (and massive changes in gene transcription) in healthy animals (**Fig. 8Ciii**).

We observed no overt differences in astrocytes in Kbk relative to wild type mice. Astrocytes are known to be triggered by activated microglia^65^. The effects of TCF-induced development non-diseased surveillance microglia on astrocytes needs investigation, but since TCF-microglia present non-inflammatory signatures, they are not likely to trigger activation of neurotoxic astrocytes^65^. Although TCF is sufficient to elicit robust transcriptional responses in Kbk microglia, it failed to stimulate post-natal neurogenesis. Prior studies showing stimulation of post-natal hippocampal neurogenesis by the HDACi AR-42, in Kbk mice^30^, maybe a consequence of sustained exposure after daily administration (at 10 mg/kg; that had to be discontinued after two weeks due to toxicity). In contrast, TCF (resulting in a single shot of 2-4 h of low micromolar concentration of Vo in the brain on a weekly basis) is well tolerated for many months in both mice and rats^22^ ^23, 66^. The improved cognitive and tactile responses in KS mice, without boosting post-natal neurogenesis suggest a role for microglia in hippocampal brain functions in Kbk mice and as targets for new therapies for KS as well as other chromatin disorders linked to intellectual disability.

## Online Materials and Methods

### Materials

All fine chemicals were purchased from Sigma (St Louis, MO, USA) unless otherwise indicated. Vorinostat was procured from Selleck Chemicals (Houston, TX, USA).

### Animals

Breeding pair of kabuki mouse model *Kmt2d^+/βGeo^* (cataloged as *Kmt2d^Gt(RRT024)Byg^*) were obtained from Bay Genomics, University of California. These mice carried a heterozygous mutation in *Kmt2d* gene and were on the mixed background of C57BL/6J and 129/SvEv. The *Kmt2d^+/βGeo^* mice were backcrossed with WT C57BL/6J mice in Haldar lab and strain purity (99-100%) confirmed by genetic analysis at Jackson labs. Breeding was performed in-house by pairing female wild type (WT) mice with male heterozygous mutant mice. Genotyping of the progenies was performed using PCR. WT allele was identified using primers, forward, 5’-CCGAGTACCTGAAAGGCGAA-3’and reverse, 5’-CTAATCAGGGCTAAGGGCGA-3’ whereas mutant allele was identified by forward, 5’-GTGTGGAACCGCATCATTGAG-3’ and reverse, 5’-GTCTTTGAGCACCAGAGGACA-3’. On agarose gels, one single band of size 411 bp suggested WT, whereas *Kmt2d^+/^*^β*Geo*^ mice produced two bands of sizes 411 bp and 272 bp.

### Drug injections and organ harvest

The Triple combination formulation (TCF) was a mixture of vorinostat (50mg/Kg), 2 hydroxypropyl-*β*-cyclodextrin (HPBCD, 2000mg/Kg), polyethylene glycol 400 (PEG, 45%) and DMSO (5%). Vehicle contained HPBCD (2000 mg/Kg), PEG 400 (45%) and DMSO (5%). Vorinostat (50 mg/Kg) was prepared in 5% DMSO and 45% PEG. Detailed methodology for preparing TCF drug solutions has been described earlier ^22, 23^. The injection volume was 10ml/Kg. Throughout the experiment, TCF or its components were administered weekly through the intraperitoneal route, post-weaning starting at 21-23 days of age. Mice were sacrificed by asphyxiation using CO_2_. The brain was isolated, the hemispheres were separated, flash-frozen in liquid nitrogen and stored at −80 °C or immersion fixed in 10% neutral buffered formalin.

### RNA extraction and qPCR analysis

Total RNA from frozen brain hemisphere was extracted using RNeasy Plus Universal kit from Qiagen and quantitative PCR (qPCR) was performed using Power SYBR Green RNA-to-CT 1-Step Kit. A 131 bp transcript region corresponding to exon 52 of *Kmt2d* (region deleted in one of the alleles in mutant mice) was detected using primers, forward, 5′-GCACCTTCACAGGCGAGACC-3′ and reverse, 5′-GAGGCCCTGGATACGCGAAC-3′. Endogenous control was *Gapdh* (*Glyceraldehyde 3-phosphate dehydrogenase*) and amplified using primers, forward, 5′-TCCATGACAACTTTGGCATTG-3′ and reverse, 5′-CAGTCTTCTGGGTGGCAGTGA-3′. The relative quantification of gene expression was done by the *ΔΔ*C_T_ method.

### Histone extraction and western blotting

Frozen brain was homogenized in Dounce homogenizer, and the histones were extracted using ‘EpiQuik Total Histone extraction kit’ (Epigentek, NY, USA) as per manufacturer’s instructions. Antibodies to acetylated histone H3, K14 (Catalog# 7627) and trimethylated histone H3K4 (Catalog# 9727) were from Cell Signaling Technology. Appropriate HRP-conjugated secondary antibodies were from Bio-Rad (Hercules, CA, USA). Quantification of band intensity was performed using ImageJ software (NIH, MD, USA).

### Isolation of hippocampus and single cell preparations

The study included a total of 60 mice, either WT or Kbk, at either one month or three-four months of age with all cohorts of mice balanced between both sexes. The one-month-old group consisted of 12 untreated mice (2 genotypes × 2 genders × 3 mice per gender). The 3-4-month-old group was divided into four treatment categories: untreated, PEG [dimethyl sulfoxide, 5% (DMSO) and polyethylene glycol, 45% (PEG)], HPBCD [DMSO (5%), PEG (45%), Hydroxypropyl-β-cyclodextrin, 0.2 gm/ml (HPBCD)], and TCF [DMSO (5%), PEG (45%), HPBCD (0.2 gm/ml), Vorinostat (5 mg/ml, Vo)]. The 3–4-month-old cohort comprised 48 mice (2 genotypes (WT and Kbk) × 2 genders (males and females) × 4 treatments (Untreated, PEG(+ DMSO), HPBCD(in +DMSO+PEG), TCF) × 3 mice per gender), Due to the large sample size, the experiment was conducted in batches (for more information, see **Fig S2A**).

Flowcharts outlining the steps involved in single-cell isolation, processing, and analysis are summarized in Figures S2B and S2C, Established methods^67, 68^, were used to generate single-cell suspensions from hippocampus tissue. Briefly, mice were anesthetized with carbon dioxide and perfused using fresh, ice-cold perfusion buffer (PB+), which consisted of Hibernate A-minus Calcium (HA-Ca; BrainBits), Actinomycin D (5 µg/ml; Sigma-Aldrich; A1410), and Triptolide (10 µM; Sigma-Aldrich T3652). The brain was carefully harvested, placed in a 10 cm dish with cold PB+, and the hippocampi were dissected out and transferred into the buffer. Using a razor blade, the hippocampi were minced into fine pieces, transferred to a 15 mL tube with PB+ using a 1 mL pipette tip with blunt end, and tissue pieces were allowed to settle. The PB+ supernatant was discarded, and the minced tissue was digested in 5 mL activated papain dissolved in dissociation buffer (DXB+) supplemented with Anisomycin (27.1 µg/ml; Sigma-Aldrich; A9789) and DNase I (2 mg/ml; Roche 10104159001). Activated papain was prepared by reconstituting lyophilized papain with 5 mL DXB+, incubating at 37°C for 10 minutes, and adding 100 µL DNase I (2 mg/ml). The minced tissue was incubated with the activated papain at 37°C for 30 minutes with gentle shaking. Afterward, the tissue was triturated initially using 5 mL pipette followed by 1 mL pipettes (10 strokes each) until no clumps remained. An additional 200 µL DNase I (2 mg/mL) was added, and the mixture was centrifuged at 300g for 5 minutes at RT.

The resulting pellet was resuspended in an ovomucoid solution (2.7 mL DXB+ with 300 µL reconstituted ovomucoid and 100 µL DNase I) to inactivate papain. The cells were layered onto 5 mL of ovomucoid solution in a separate tube and centrifuged at 300g for 10 minutes. The supernatant was discarded, and the pellet was resuspended in Hibernate A-minus Calcium (HA-Ca). The suspension was passed through a pre-wetted 70 µm filter (Miltenyi) to remove debris, and the filter was rinsed with 2 mL HA-Ca to collect any remaining cells. The final cell suspension was centrifuged at 300g for 10 minutes at 4°C (with maximum acceleration and half brake) and the cell pellet was process for debris removal as described below.

### Percoll gradient for debris removal ^69^. ^69^

The pellet was resuspended in 1 ml 30% Percoll in PBS (1X, pH 7.4) and carefully layered on 5 ml 70% Percoll (made in PBS (1X, pH 7.4) and another 4 ml of 30% Percoll was layered carefully on the top of cell suspension. The gradient was subjected to centrifugation for 30 minutes at 300g at 22°C. Debris-free single cells were collected from the 70%-30% Percoll interface and transferred using a 1 ml pipette with blunted tip to another 15 ml tube containing 14 ml Hibernate A-Low Fluorescence (HA-LF) containing Actinomycin, Triptolide, and Anisomycin. The mixture was centrifuged for 10 min, 300*g*, 4°C, with maximum acceleration and brake. The cell pellet was resuspended in 300 µL of HA-LF and corresponding inhibitors.

### Cell Hashing protocol^70^

For cell hashing, the cells were counted using a hemocytometer. One million cells were resuspended in 100 µL of Staining buffer (2% BSA/0.02% Tween, PBS), blocked with 10 µL of Fc Blocking reagent (BioLegend), and incubated at 4°C for 10 minutes. After blocking, 1 µg of hashing antibody was added, and the mixture was incubated for 30 minutes at 4°C. The cells were then washed three times with Staining buffer, pelleted at 400 × g for 5 minutes at 4°C, and resuspended in 1X PBS, pH 7.4.

### Single cell capture, library preparation and sequencing

The percent viability and concentration of the cell suspension was determined by staining Trypan Blue Stain (0.4%; ThermoFisher Scientific PN-T10282) and Countess II Automated Cell Countess. It was loaded into Chromium Next GEM Chip G (10X Genomics PN-1000127) with a targeted recovery of 16,000 to 20,000 cells per well and processed with Chromium Controller (10X Genomics). Single-cell RNAseq libraries were prepared with Chromium Single Cell 3’ Reagent Kit (v3.1 CG000204 Rev C; 10X Genomics PN-1000128) and Chromium i7 Multiplex Kit (Single Index; PN-120262) following the standard throughput manual workflow. Single-cell HTO libraries were prepared following CITE-seq & Cell Hashing Protocol (New York Genome Center Technology Innovation Lab, version 2019-02-13). The single-cell RNA-seq libraries were sequenced on an Illumina NovaSeq 6000 with paired-end reads (28 cycles Read 1, 8 cycles i7, 91 cycles Read 2).

### Single cell data analysis

The Cell Ranger software (10x Genomics) was used to align the FASTQ sequencing data from the individual batches to the reference mouse transcriptome (using STAR algorithm) and the demultiplexing was performed by unique molecular identifiers (UMI) and cell specific barcodes. After demultiplexing, the number of unique molecular identifiers (UMIs) associated with each gene in each cell was quantified by Cell Ranger. The Cell Ranger output consists of three matrices where rows of feature-barcode matrix represent genes, columns represent cells, and each entry represents the expression level of a gene in a specific cell. The filtered expression matrix output of each batch was processed by R package Seurat_5.1.0. The data from each batch was (total eight batches) merged into a single Seurat object and subsequently, the single cells were subset and used for further analysis. The integrity of individual cells obtained was assessed by filtering out unhealthy and dying cells, after considering the number of genes detected, their counts and the proportion of mitochondrial genes. The cells with transcripts number below 500 or exceeding 8000 and transcript counts exceeding 30000 as well as those with a mitochondrial gene percentage more than 10% per cell, were excluded from further analysis (**Fig S2D**).

### Normalization, highly variable gene selection, scaling, and dimensionality reduction

After filtering out unhealthy cells we normalized the dataset by log normalization. The log-transformed values are then centered across all genes for each cell, resulting in the relative expression levels across different cells. Next, we selected 3000 highly variable genes which are highly expressed in some sets of cells and at very low levels in other sets of cells. These genes have significantly higher variance across cells than by chance. The genes with high variance are likely to be more informative and may represent the biologically meaningful differences between cell types or states. After highly variable genes are selected, to negate the effect of their high expression difference, data is scaled to bring the expression levels of genes within the range. Scaling helps to remove the unnecessary source of variation of the data by bringing the distribution of values between 0 and 1. To reduce the dimensionality of the large matrix to small matrix with less factors, principal component analysis (PCA) was performed using the first 50 PCAs and highly variable genes selected (which were scaled in the previous step) were used for the PCA input. The PCA projects a high dimensional dataset to very few dimensions. The top few principal components capture the majority of variance of the data are kept for the downstream steps.

### Batch correction, cell clustering and cell type identification

The PCA transformed data is fed into the batch correction step by harmony method ^71^ using R package harmony_1.2.0. It is fast and removes technical variations introduced by batch effects, while preserving biological variability across batches and experimental conditions. Next, to visualize the highly complex data, we generated UMAP using harmony embeddings. Cell clustering and cell type identification was done with the batch corrected data. Seurat uses graph-based clustering method. The shared nearest neighbor (SNN) identifies cell to cell relationships based on the nearest neighbor graph to define cell neighborhoods. To cluster the cells, Louvain algorithm^72^ was used iteratively to group cells together. It tries to determine the modularity of the graph. In network analysis modularity is a commonly used concept to identify the community structure within a graph. It measures how well communities can be partitioned into distinct structures and suggest a strong community. We used a resolution parameter of 0.6 while clustering and resulted in 28 clusters (0-27) overall. We used canonical cell type markers to identify the distinct brain cell types (Table S2, Fig. S2E).

### Differential gene expression and heatmaps of microglia subset

The differential gene expression analysis was performed on microglial subset using R package DESeq2_1.44.0 ^73^. Since the count dispersion of features across samples is quite high, and to account for this overdispersion in scRNAseq data, DESeq2 uses negative binomial model to obtain a better statistical significance measure. DESeq2 uses the Wald test for pair wise analysis between groups. The raw p-values were calculated using the Wald test and is a test of hypothesis usually performed on parameters that have been estimated by maximum likelihood. The significant genes filtered both by raw p-values (<0.05) and adjusted p-value (padj).

### Trajectory analysis

The preprocessed, integrated, and batch corrected microglia single cell RNA sequencing data (microglia Seurat object) was used as the input for trajectory analysis. Trajectory analysis was performed using the monocle3_1.3.7 package in R to understand the developmental trajectory of single cells. Initially, the Seurat object is converted to cell data set object (cds) and the cds object was further pre-processed by following steps. The cds were preprocessed using the preprocess_cds() function, which normalized the data using principal component analysis (PCA) with the first 50 dimensions. UMAP was used to reduce dimensionality to two axes. A trajectory graph was constructed by learn_graph() function. Once the trajectory graph was constructed, the cells were ordered along the trajectory depending on their progress through the development program. We selected one-month untreated wild type group (WT.1m.Un) as the root node in the current analysis since this time point was only a week after the pups were released from weaning. The trajectory graph was visualized using UMAP plot. The pseudotime values generated for all the 12216 cells and stored in the cds. The pseudotime vector stored in the cds was extracted and added to the metadata of microglia Seurat object. The continuous pseudotime vector was further converted to 25 categories (25 bins) and the resulting vector was stored as a new variable “pseudotime_bin” in the metadata of microglia Seurat object. Next, we calculated the proportion of each group in the bins and plotted it as a trajectory vs proportion graph.

### Module score analysis

To assess the activity of specific gene sets in individual cells, we performed module score analysis using AddModuleScore function of the Seurat_5.1.0 package in R. Module scores were calculated for Homeostatic, Stage 1 DAM and Stage 2 DAM gene sets. After splitting the microglia Seurat object based on group variable (there are 10 groups), the module score was calculated for each group from individual cells. It calculates a score for each cell representing how strongly a set of modules (set of genes), are expressed within a cell, compared to a set of randomly selected background control genes. The control genes were chosen with similar expression levels to ensure that the module score is not confounded by overall gene expression differences between cells. As data may not follow a normal distribution, the Wilcoxon rank sum test is used as default test to compare module genes to control genes.

### Novel Object Recognition assay

The object recognition test was used to assess memory in mice. The assay was conducted on three consecutive days. On day 1, mice were individually placed into a plastic container with black walls of size 37×29×17 cm and allowed to explore the arena for 10 minutes. On day 2, two identical objects were placed in the center (kept at equidistance from the wall and each other) and the time interacting with each object was recorded over 10 minutes. On the third day, one object was removed and replaced by a novel object. Lego towers and T25 tissue culture flask filled with white gravel were used as objects. Mice were placed in the arena for ten minutes and timed for interaction with each object. The arena and objects were cleaned with 70% ethanol between each mouse to eliminate olfactory cues. The discrimination index was calculated by using a formula, TN-TF/TN+TF, where TN and TF are time spent exploring novel and familiar objects, respectively.

### Von Frey Assay

Tactile sensitivity is measured by studying paw withdrawal frequency (PWF) using a series of von Frey filaments with incremental filament thickness ^58^. The mice are placed into an acrylic box and habituated in a quiet room for an hour. Mechanical stimulus applied perpendicularly to the mid-plantar surface of hind paws for 2 seconds with enough force to bend the filament. Each paw was stimulated 5 times by each filament with an interval of 5 seconds between each measurement. Only rapid paw withdrawal or prolonged paw withdrawal combined with flinching or licking of the paw were recorded. 50% paw withdrawal threshold was calculated as described ^58^. The left and right hind paws of each mouse were assessed on three consecutive days and the average withdrawal threshold of each paw was used to plot the data. A lower 50% threshold value suggests higher tactile sensitivity.

### Immunofluorescence assay and microscopy

The brain hemisphere was immersion fixed in 10% neutral buffered formalin (∼4% paraformaldehyde) for 20-22 h and paraffin-embedded. 5µm thick sagittal sections spaced at 40µm collected at 4-6 different depths were processed for DCX staining. After dewaxing and hydration, DCX antigen was retrieved by boiling the sections in acidic condition for 30 min followed by a 20-minute incubation at room temperature as the sections cool off. After blocking in 2% goat serum, sections were incubated with anti-DCX antibody (Santa Cruz Biotechnology, sc-271390, 1:50) overnight at 4 °C. FITC conjugated secondary IgG (MP Biomedicals) was used at 1:1000 dilution. Sections were mounted using Vectashield (Vector Laboratories) containing DAPI for staining nuclei and visualized by fluorescence microscopy. DCX labeled brain sections were visualized by Olympus IX inverted fluorescence microscope using 40x oil-immersion objective lens (NA1.35) and the number of DCX+ cells in the inner margin of entire SGZ was manually counted. Digital images of consecutive 20 optical sections of 0.2 µm thickness were collected by Photometrix cooled CCD camera (CH350/LCCD) driven by DeltaVision software from Applied Precision (Seattle, WA, USA). DeltaVision software (softWoRx) was used to deconvolve these images. The images presented are maximum intensity projections of all 20 optical sections. Images were analyzed using ‘softWoRx’ or ‘ImageJ’ software (NIH, MD, USA).

### Immunohistochemistry, image acquisition and analysis

Tissue embedding and immunohistochemical staining of sagittal brain sections was performed by NeuroScience Associates (Knoxville, TN, USA). Thirty-micron thick sections were cut through the entire depth of the brain. Free-floating sections after blocking were incubated in rabbit anti IBA-1 antibody (Abcam, ab178846, 1:75000) or anti GFAP antibody (Dako, Z0334, 1:14000) for overnight at room temperature. Biotinylated secondary antibodies were from Vector Laboratories (BA-1000, 1:1000). Sections were developed following standard methods. To quantify microglial abundance in hippocampus, IBA1 stained brain slices from 6-7 depths at the interval of 180µm were analyzed and bright field images were captured using A10 PL 10x (for counting) objective lens (Nikon) with a numeric capture of 0.25. After applying constant threshold numbers were automatically counted using ImageJ. For analyzing astrocytes, 4-5 random images of GFAP-stained hippocampus at 3 depths spaced at 180µm from each mouse were captured using 40x oil-immersion objective lens (Plan-Apochromat, Nikon) with a numeric aperture of 1.3. Both objective lenses (10x and 40x) were connected to the Nikon Olympus microscope supported by Nikon digital DS-Fi1-U2 camera controlled by NIS-Elements F3.0 Nikon software. GFAP positive areas were highlighted by applying constant threshold and percentage-stained areas were quantified using ImageJ.

### Morphometric assessments of microglia using Imaris

For analyzing microglia extensions and soma, 2 random images of hippocampus (stained for IBA1 as described above) at 2 depths (4 images/mouse) spaced at 180µm were acquired using 40x oil-immersion objective lens as described above. For morphometric measurements of microglia extensions and soma, images were processed using the Imaris 9.5.1 algorithm (Bitplane, UK). All measurements were done on 2-dimensional images after converting bright filed images to ‘ims’ files. Structural complexity of extensions was analyzed by quantifying cumulative length of filaments, number of branches and terminal points. Extensions originating from soma were identified and automated tracing was performed by ‘auto path’ feature in filament wizard. The origin of filaments from the soma of single microglia was individually verified. Microglia present at the edges or soma touching each other were excluded from the analysis. Microglia with intermingled extensions or filament with incorrect connections (seen in less than 5%) were manually removed. For extensions, 400-500 hippocampal microglia (40±10 per mouse) per experimental group were reconstructed whereas cell soma was identified via surface creation wizard in 600-700 (60±15 per mouse) microglia. For soma, a constant threshold was set to exclude branches and extensions.

### Statistical Test

Graphs were plotted in MS Excel or GraphPad Prism 8.4.3. For comparing two groups student’s *t-test* or Mann-Whitney U test was performed. For comparing more than two groups, unless specified, one-way ANOVA with Tukey’s posthoc analysis (parametric) or Kruskal-Wallis test (nonparametric) was performed. All statistical tests were done with GraphPad Prism 8.4.3 and were two-tailed. Statistical test and P values are indicated in figure legends. *P<0.05* was considered significant. *\*P<0.05, **P<0.01, ***P<0.001, ****P<0.0001*.

## Acknowledgments

We thank Dr. Siyuan Zhang (University of Notre Dame) and members of the Haldar lab for helpful discussion during the work. We thank Katharyn Hutson and Vy Sanders for assisting with mouse studies.

We thank the Freimann Life Sciences Center (FLSC) and staff at the University of Notre Dame, for providing excellent services for mouse housing, caring, breeding, blood draws overall monitoring and veterinarian support.

We thank the Genomics and Bioinformatics Core Facility at the University of Notre Dame for their assistance with single cell sequencing studies.

## Regulatory Approval

All procedures were approved by the Institutional Biosafety Committee (IBC), University of Notre Dame. Animals studies and procedures regarding their use were approved by the Institutional Animal Care and Use Committee (IACUC) University of Notre Dame.

## Author Contribution

Md. Suhail Alam –conceptualization, design, experimentation, analysis, and interpretation of Figures 1, 3, 4, 6, 7, 8 (and other associated supplementary data figures), genotyping strategy and assay development for mouse colonies; organ harvest; tissue analyses; visualization of results, primary drafting and editing the manuscript.

Prasad Padmanabhan – conceptualization, design, and analyses of Figures 2, 3, 5 (and other associated supplementary data figures and tables) related to single cell rna sequencing studies, data deposition, colony management organ harvest, visualization of results, writing of related methods and figure legends as well as primary drafting and editing of the manuscript.

Ryan McArdle - conceptualization, design, and analyses of Figure 2 (and other associated supplementary data figures and tables), mouse genotyping colony management, organ harvesting, visualization of results, and editing of the manuscript.

Alejandro Lopez-Ramirez - conceptualization, design, and analyses of Figures 3, 4, 10 (and associated supplementary data), behavioral assays, mouse genotyping, organ harvesting visualization of results and editing of the manuscript.

Arpitha MysoreRajashekara – mouse genotyping, behavioral assays, colony management, organ harvest, visualization, data analyses and editing of the manuscript.

James Knopp-quantitative analyses and data visualization of hippocampal microglia, morphometric and activation characteristics using Imaris.

Gabriela Kim - quantitative analyses of hippocampal microglia using ImageJ.

Maria Virginia Centeno – training and analysis of von Frey test for tactile sensitivity in rodents. Apkar Vania Apkarian – data analyses, interpretation and overall expertise in nociception and pain and editing the manuscript.

Vivek Swaroop – overall conceptualization, design, and project supervision of single cell RNA sequencing studies, primary drafting and editing of the manuscript.

Kasturi Haldar – overall conceptualization and project supervision, design, and development of mouse *in vivo* and *in vitro* studies; visualization of results; primary drafting and editing of manuscript, funding acquisition.

All authors reviewed and commented on the manuscript and approved of its contents.

