## Supplementary Figures for "Chronic HDACi emends microglial differentiation and neurological disease in a mouse model of intellectual disability"

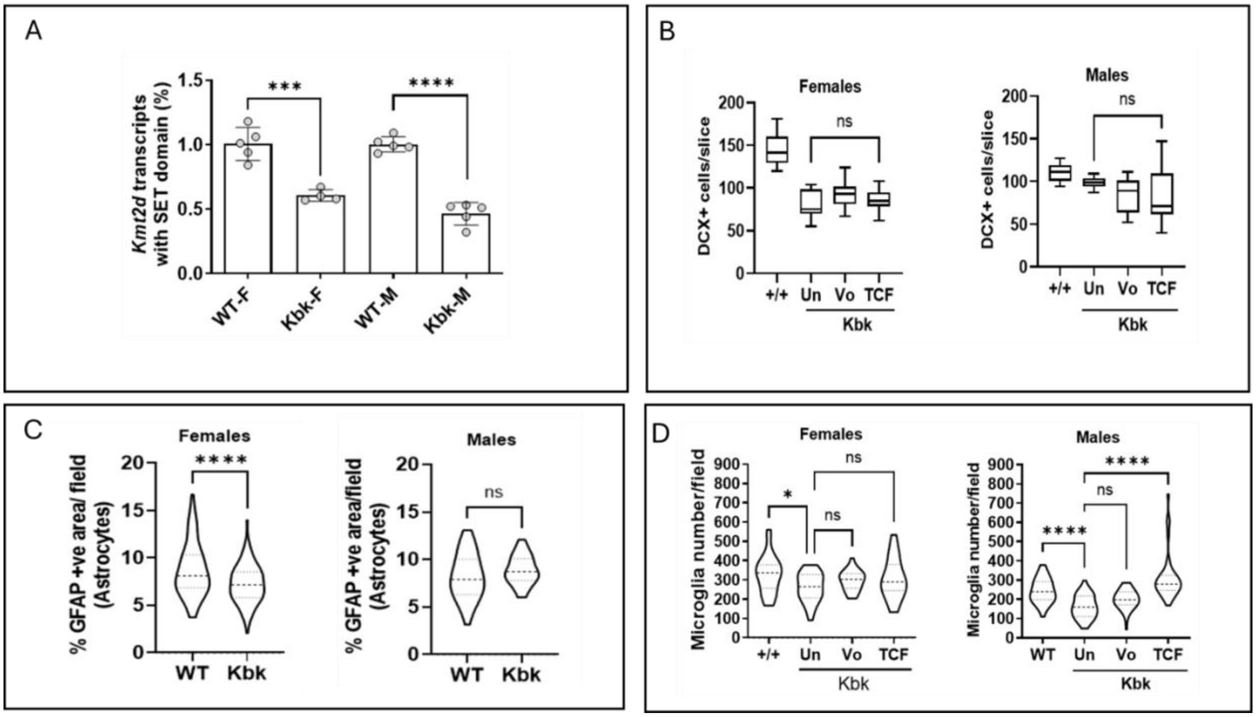


**Supplementary Fig. S1. Effect of sex on molecular and cellular characteristics of Kbk mutant compared to wild type brain. A.** qPCR analysis of *Kmt2d* transcripts (measured by detection of exon 52) in male (M) and female (F) Kbk mice compared to wild type (WT) counterparts. Data are mean±SEM **B.** Quantification of DCX (doublecortin X) positive neuroblasts in the entire subgranular zone (SGZ) of the dentate gyrus (DG, measured over the entire depth of the hippocampus; 5-6 sections) of WT or Kbk mutants untreated (Un) or treated with vorinostat (Vo) or TCF; n=4-6 mice/group. Statistical analysis was conducted by one-way ANOVA with Tukey's posthoc test. **C.** Quantitative analysis of GFAP- labeled astrocytes in the hippocampus from WT and Kbk mice. Granule cell regions and the polymorph layer of the DG were excluded to eliminate neural progenitors (that are also labeled with GFAP). Percentage GFAP positive areas calculated in 4-5 random fields at three depths (across all mice) are shown. Statistical analysis by Mann-Whitney test. **D**. Violin plots show quantification of IBA1 positive microglia in hippocampus after no treatment (Un), treatment with vorinostat (Vo) or TCF. Statistical analyses used one-way ANOVA with Tukey's post hoc test. ns, non-significant. **P<0.05, ***P<0.001, ***P<0.0001.*

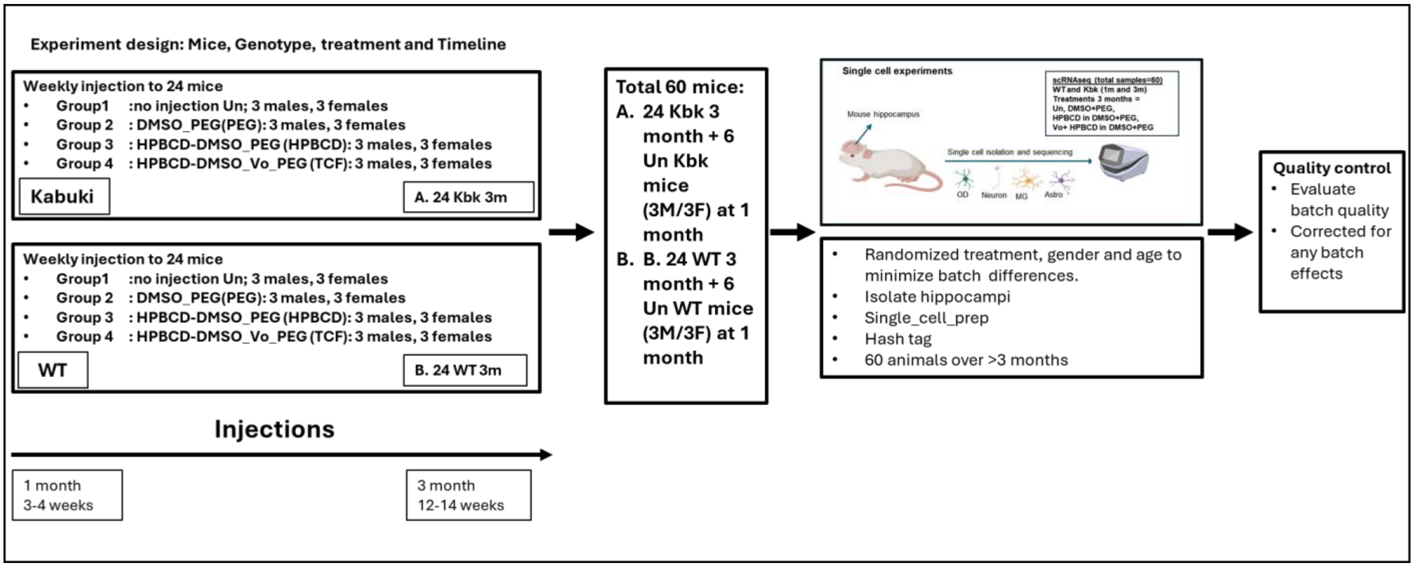


**Supplementary Fig. S2A. Summary of experimental design of single cell RNA sequencing of wild type and mutant microglia and their responses to chronic TCF and its components.** Twenty-four Kabuki (Kbk) and twenty-four wild type (WT) mice at one month of age (equal numbers of males and females) were separated into four sets of 6 (3 males, 3 females). For both Kbk and WT groups, one set was left un-injected, while the remaining three sets received weekly injections of TCF or control treatments of DMSO+PEG or HPBCD+DMSO+PEG, up to three months of age. For subsequent analyses, to both Kbk and WT groups, we added another set of six untreated (Un) mice (3 males, 3 females) at one month of age. Parameters of treatment, age and gender were randomized to minimize batch-differences. Hippocampi were purified and subsequently isolated cells were processed through the indicated single cell pipeline where the batch-quality was closely monitored to eliminate batch-effects (see Figs. S2B-S2D). Canonical microglial markers and proportion graphs were obtained as described in Figs. S2E-F.

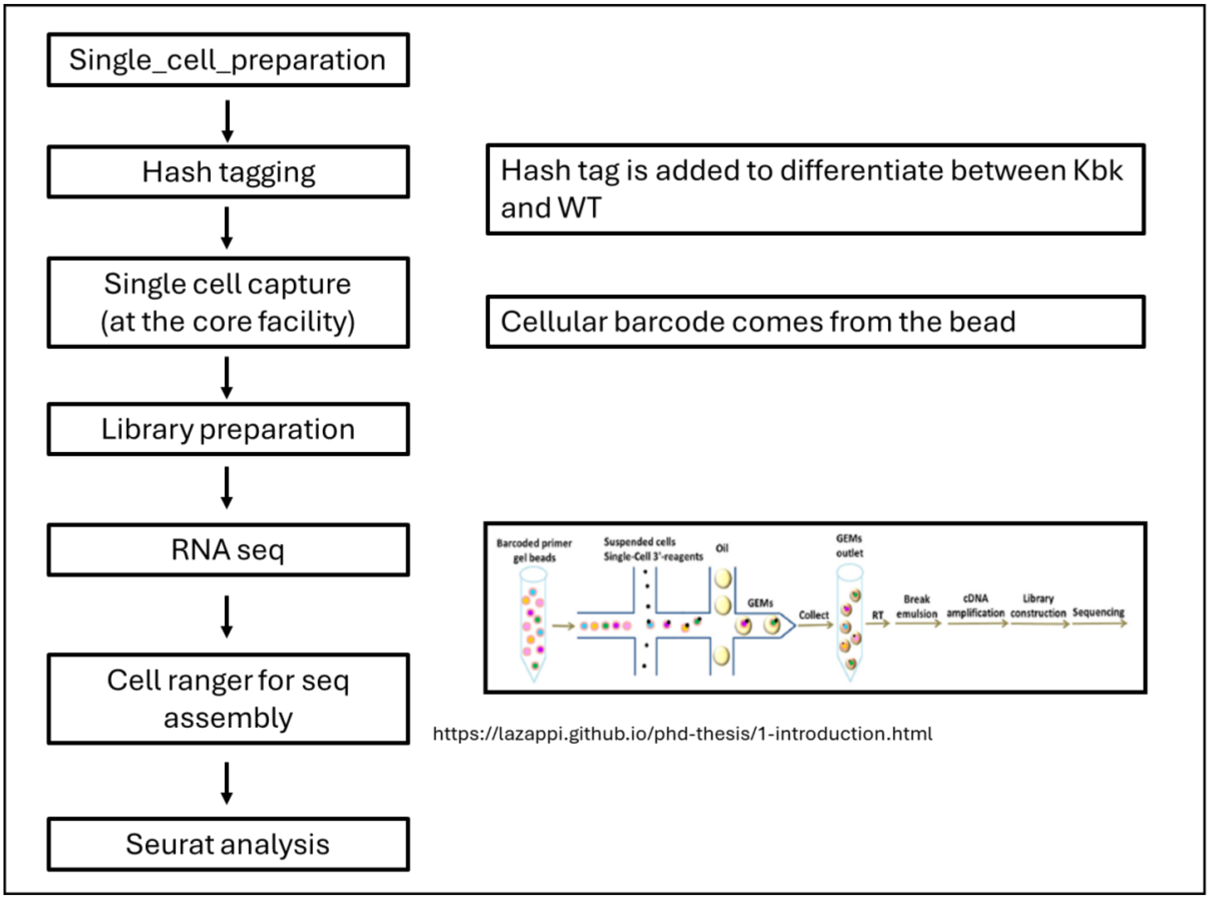


**Supplementary S2B. Overview of the main steps of single-cell RNA sequencing analysis.** Isolated cells were hash-tagged to differentiate between Kbk and WT, underwent single cell capture (by barcoded beads), library preparation, RNA seq, cell ranger for seq assembly and Seurat analyses.

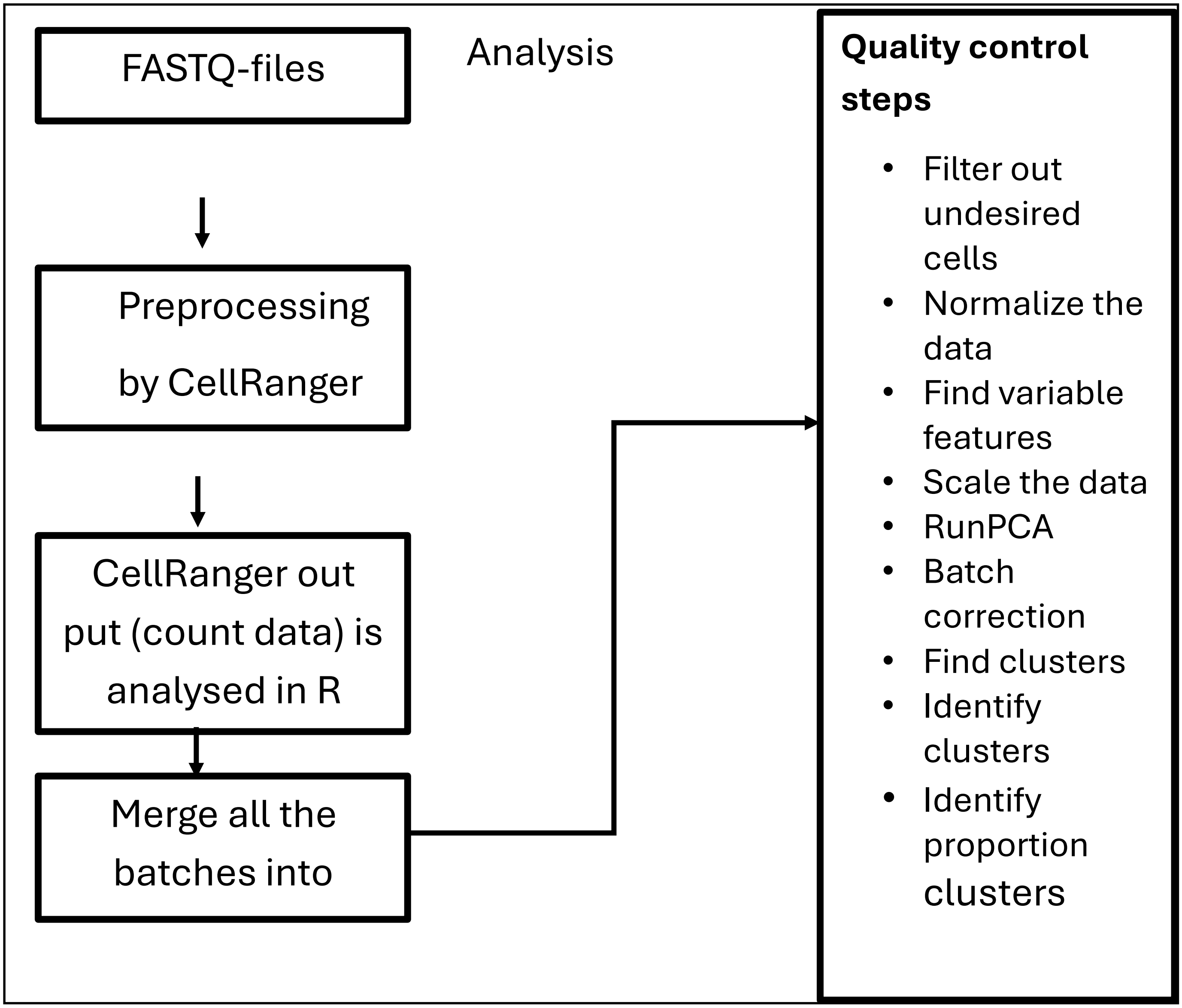


**Supplementary S2C**. **Preprocessing steps undertaken after single-cell RNA sequencing.** FASTQ sequences were inputted into the Cell Ranger pipeline, which processed them by aligning the reads to a reference genome and generated count data for each gene.

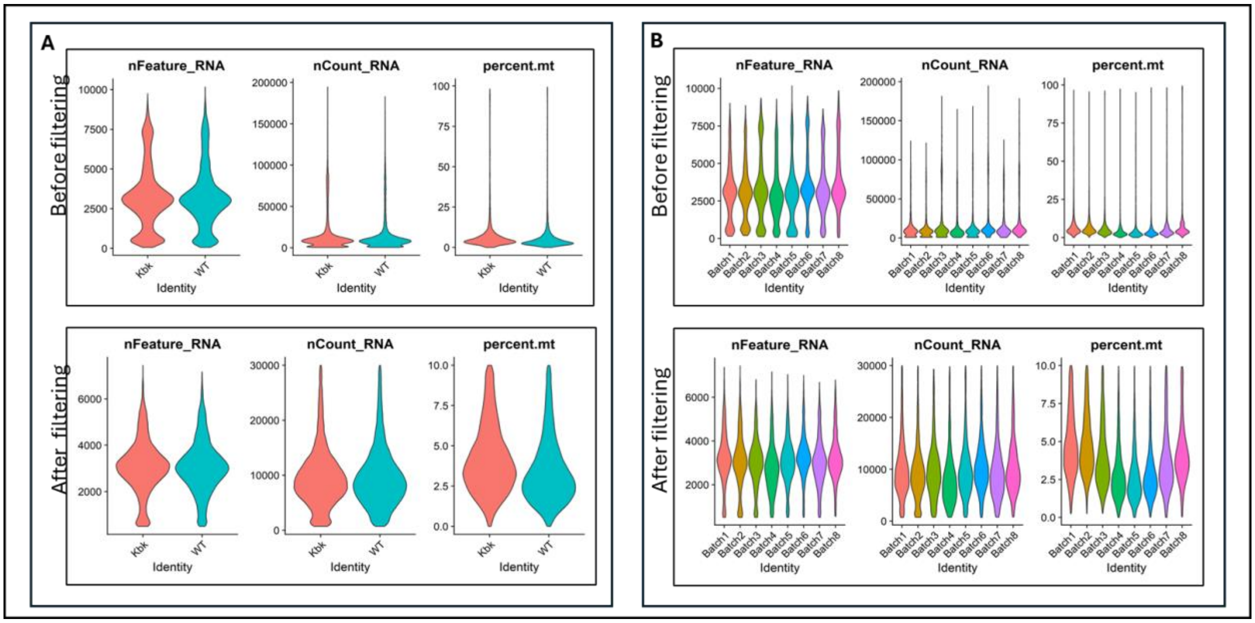


**Supplementary Fig. S2D**. **Quality control of single cell analyses**. Removal of low-quality information was done using Seurat in the following way. Cells with fewer than 500 detected genes (nFeature_RNA), more than 10% mitochondrial gene expression and transcript counts (nCount_RNA) exceeding 30000, were excluded. **A.** Distribution of genes (nFeature_RNA), gene counts (nCount_RNA) and mitochondrial percentage(percent.mt) across WT and Kbk genotypes before and after filtering for low quality cells. **B.** Distribution of genes (nFeature_RNA), gene counts (nCount_RNA) and mitochondrial percentage (percent.mt) across **batches** before and after filtration of low quality cells.

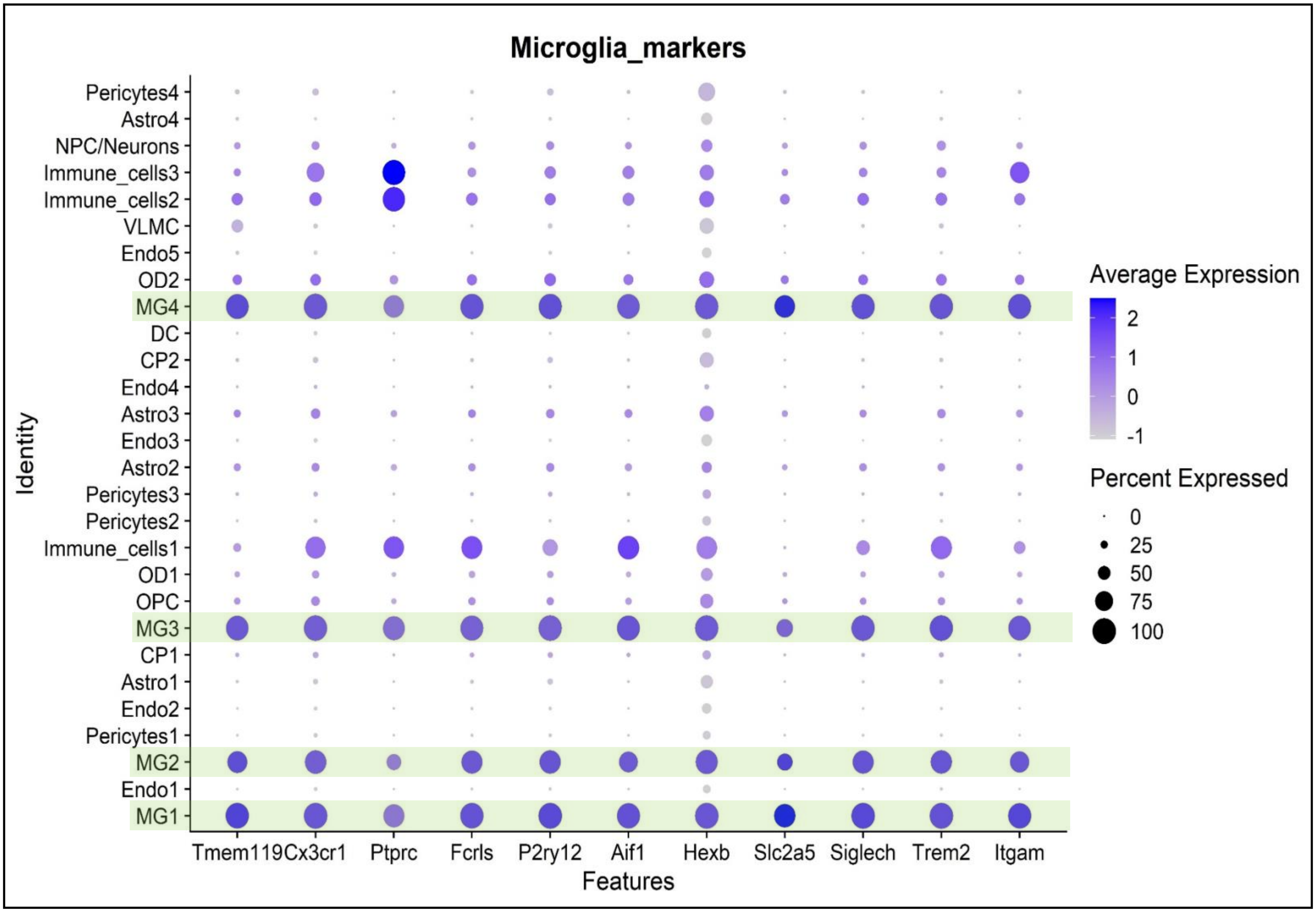


**Supplementary Fig. S2E. Dot plot representation of canonical microglial markers**. Four microglial clusters (MG1, MG2, MG3, MG4, indicated by green horizontal bars) were enriched for 11 canonical markers, as shown.

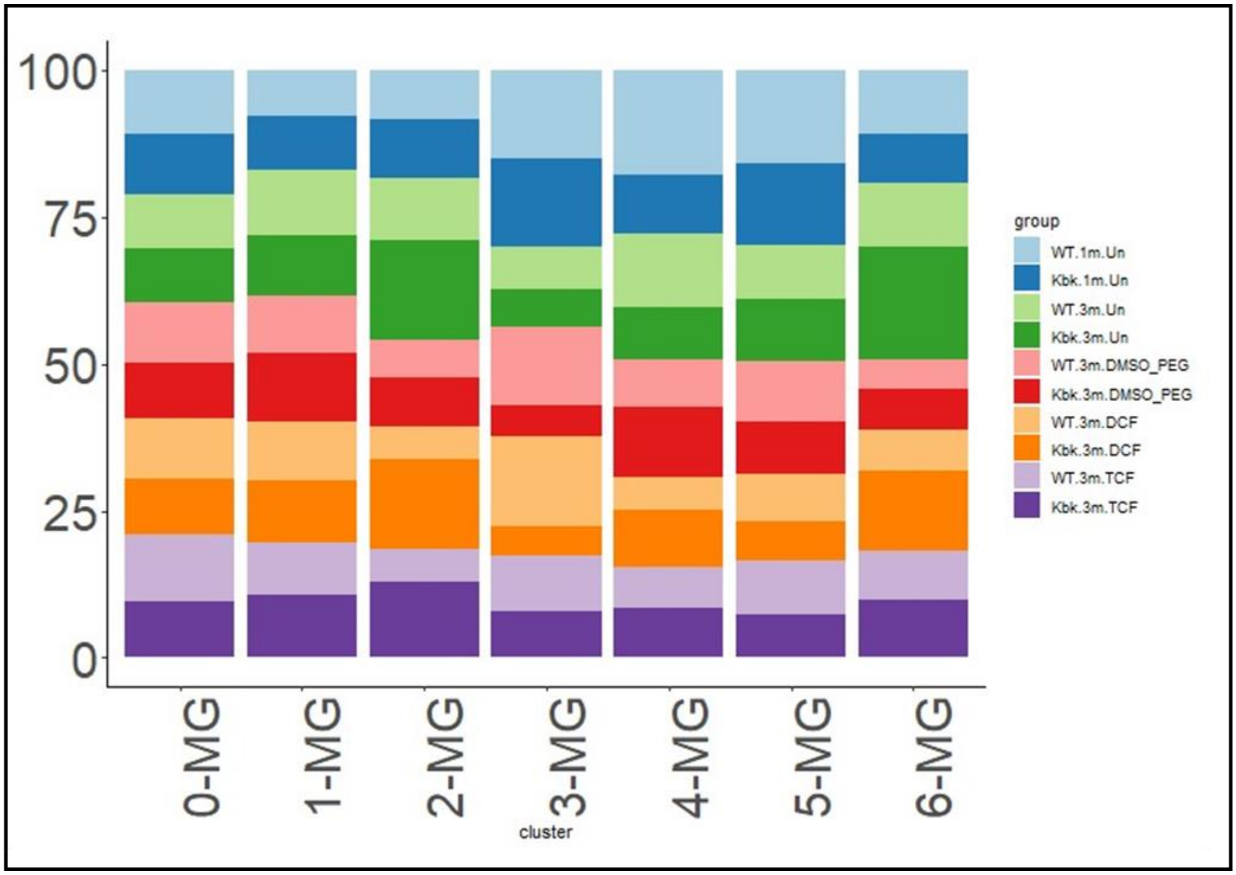


**Supplementary Fig. S2F**: **Proportion maps**. Data were extracted from original cell type clusters and re-clustered using R package Seurat. The proportion of each of the microglial subsets were calculated and visualized using ggplot2 package

**Supplementary Fig. S3. Differential gene expression in hippocampi of Kbk compared to wild type (WT) under different conditions**. **A. Differential gene expression** (determined using DESeq2 package of R) induced by HPBCD(in DMSO+PEG) compared to PEG (in DMSO) at 3 months in Kbk mice. Genes with p-value threshold of 0.05 were filtered and used for the above heat map. Asterisks correspond to genes filtered using padj (p-adjusted) <0.05. **B. Top 30 pathways changed in Kbk versus WT** at 3 months of age as predicted by IPA**. C Top 30 pathways changed as in Kbk versus WT** at one month of age, as predicted by IPA**.**

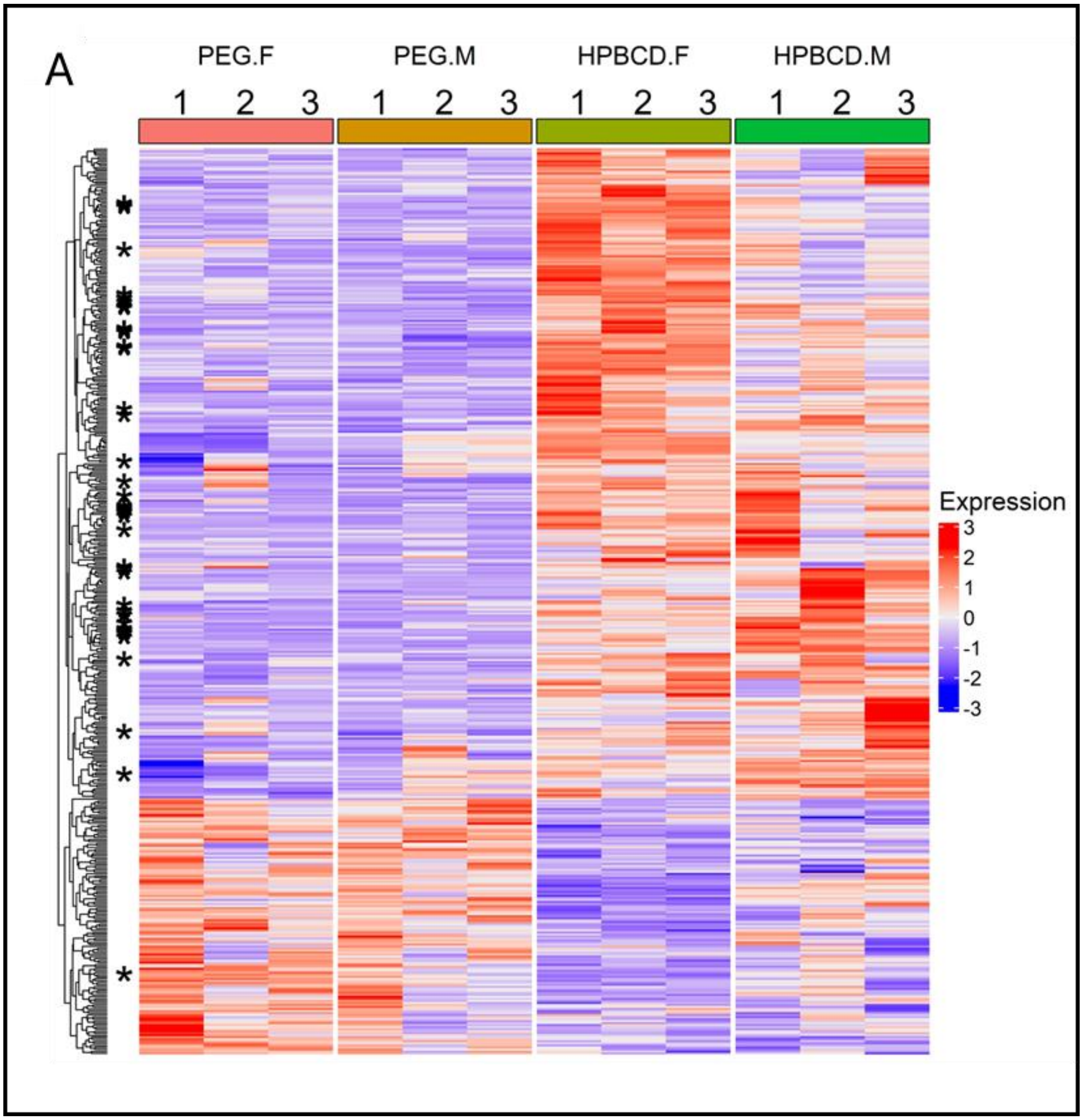

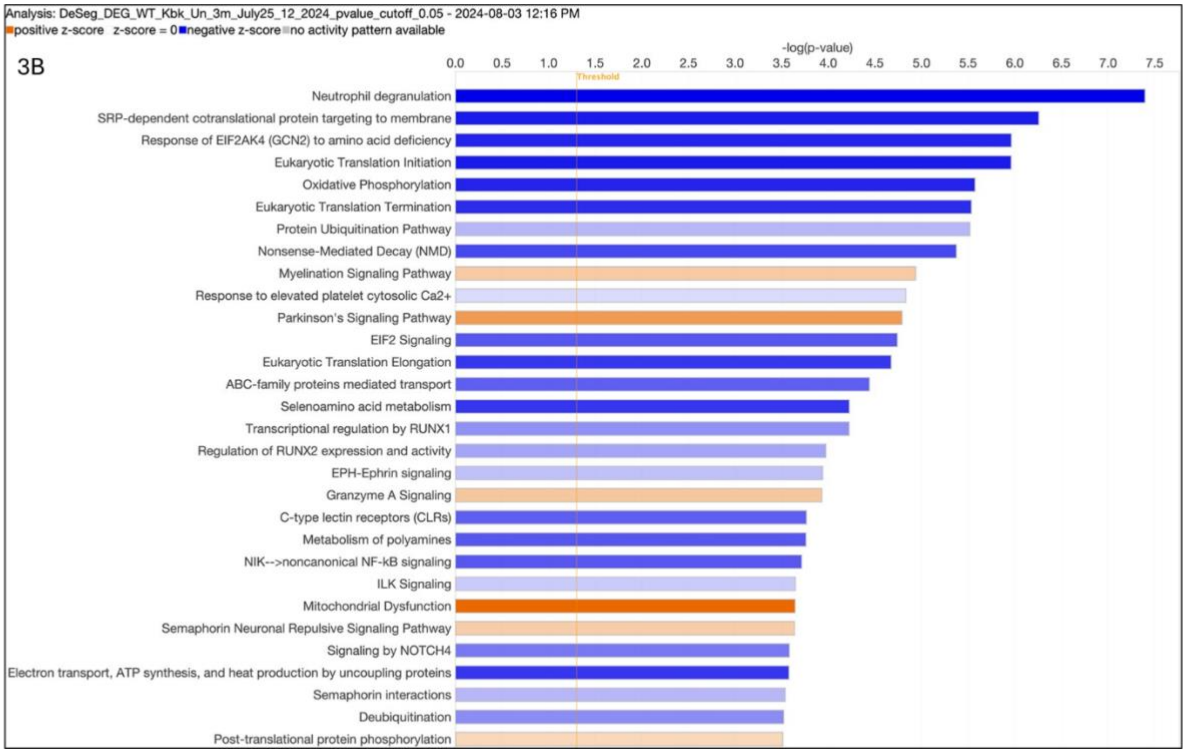

Analysis: DeSeg_DEG_WT_Kbk_Un_3m_July25_12_2024_pvalue_cutoff_0.05 - 2024-08-03 12:16 PM
positive z-score z-score = 0 negative z-score no activity pattern available
-log(p-value)
3B
0.0 0.5 1.0 1.5 2.0 2.5 3.0 3.5 4.0 4.5 5.0 5.5 6.0 6.5 7.0 7.5
Threshold
Neutrophil degranulation
SRP-dependent cotranslational protein targeting to membrane
Response of EIF2AK4 (GCN2) to amino acid deficiency
Eukaryotic Translation Initiation
Oxidative Phosphorylation
Eukaryotic Translation Termination
Protein Ubiquitination Pathway
Nonsense-Mediated Decay (NMD)
Myelination Signaling Pathway
Response to elevated platelet cytosolic Ca2+
Parkinson's Signaling Pathway
EIF2 Signaling
Eukaryotic Translation Elongation
ABC-family proteins mediated transport
Selenoamino acid metabolism
Transcriptional regulation by RUNX1
Regulation of RUNX2 expression and activity
EPH-Ephrin signaling
Granzyme A Signaling
C-type lectin receptors (CLRs)
Metabolism of polyamines
NIK-->noncanonical NF-kB signaling
ILK Signaling
Mitochondrial Dysfunction
Semaphorin Neuronal Repulsive Signaling Pathway
Signaling by NOTCH4
Electron transport, ATP synthesis, and heat production by uncoupling proteins
Semaphorin interactions
Deubiquitination
Post-translational protein phosphorylation


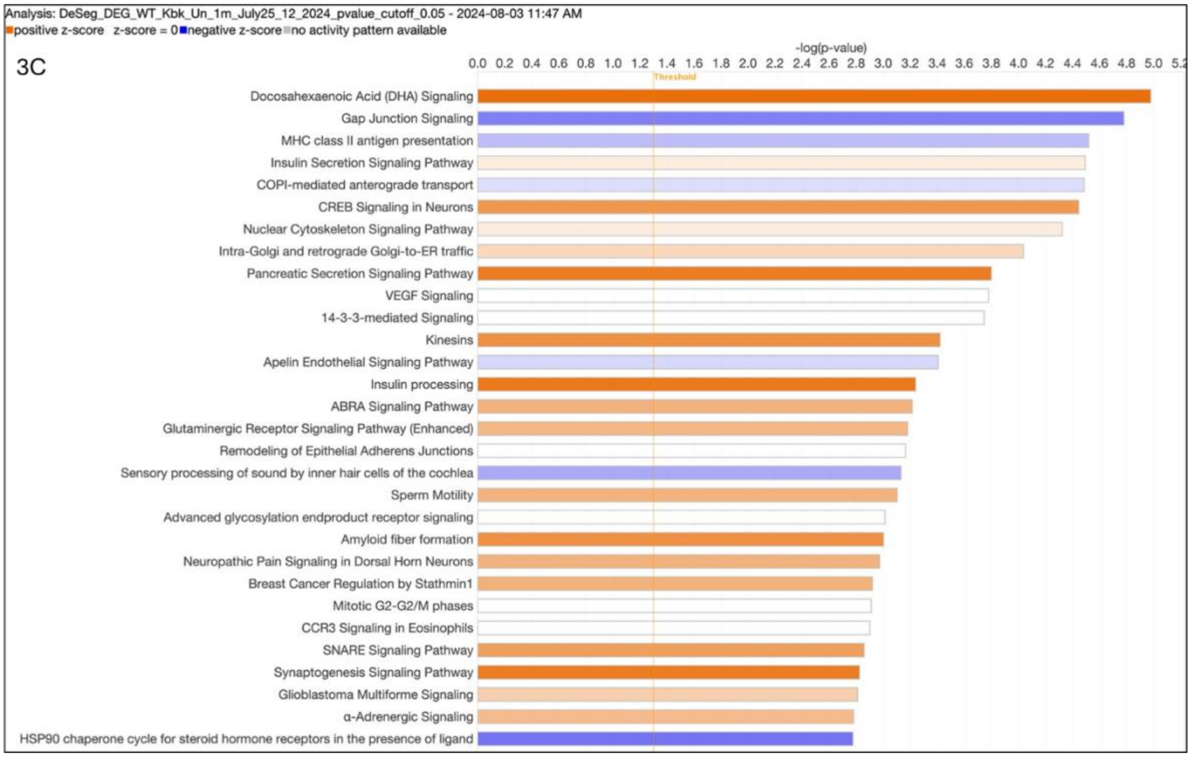

Analysis: DeSeg_DEG_WT_Kbk_Un_1m_July25_12_2024_pvalue_cutoff_0.05 - 2024-08-03 11:47 AM
positive z-score z-score = 0 negative z-score no activity pattern available
-log(p-value)
3C
0.0 0.2 0.4 0.6 0.8 1.0 1.2 1.4 1.6 1.8 2.0 2.2 2.4 2.6 2.8 3.0 3.2 3.4 3.6 3.8 4.0 4.2 4.4 4.6 4.8 5.0 5.2
Threshold
Docosahexaenoic Acid (DHA) Signaling
Gap Junction Signaling
MHC class II antigen presentation
Insulin Secretion Signaling Pathway
COPI-mediated anterograde transport
CREB Signaling in Neurons
Nuclear Cytoskeleton Signaling Pathway
Intra-Golgi and retrograde Golgi-to-ER traffic
Pancreatic Secretion Signaling Pathway
VEGF Signaling
14-3-3-mediated Signaling
Kinesins
Apelin Endothelial Signaling Pathway
Insulin processing
ABRA Signaling Pathway
Glutaminergic Receptor Signaling Pathway (Enhanced)
Remodeling of Epithelial Adherens Junctions
Sensory processing of sound by inner hair cells of the cochlea
Sperm Motility
Advanced glycosylation endproduct receptor signaling
Amyloid fiber formation
Neuropathic Pain Signaling in Dorsal Horn Neurons
Breast Cancer Regulation by Stathmin1
Mitotic G2-G2/M phases
CCR3 Signaling in Eosinophils
SNARE Signaling Pathway
Synaptogenesis Signaling Pathway
Glioblastoma Multiforme Signaling
α-Adrenergic Signaling
HSP90 chaperone cycle for steroid hormone receptors in the presence of ligand

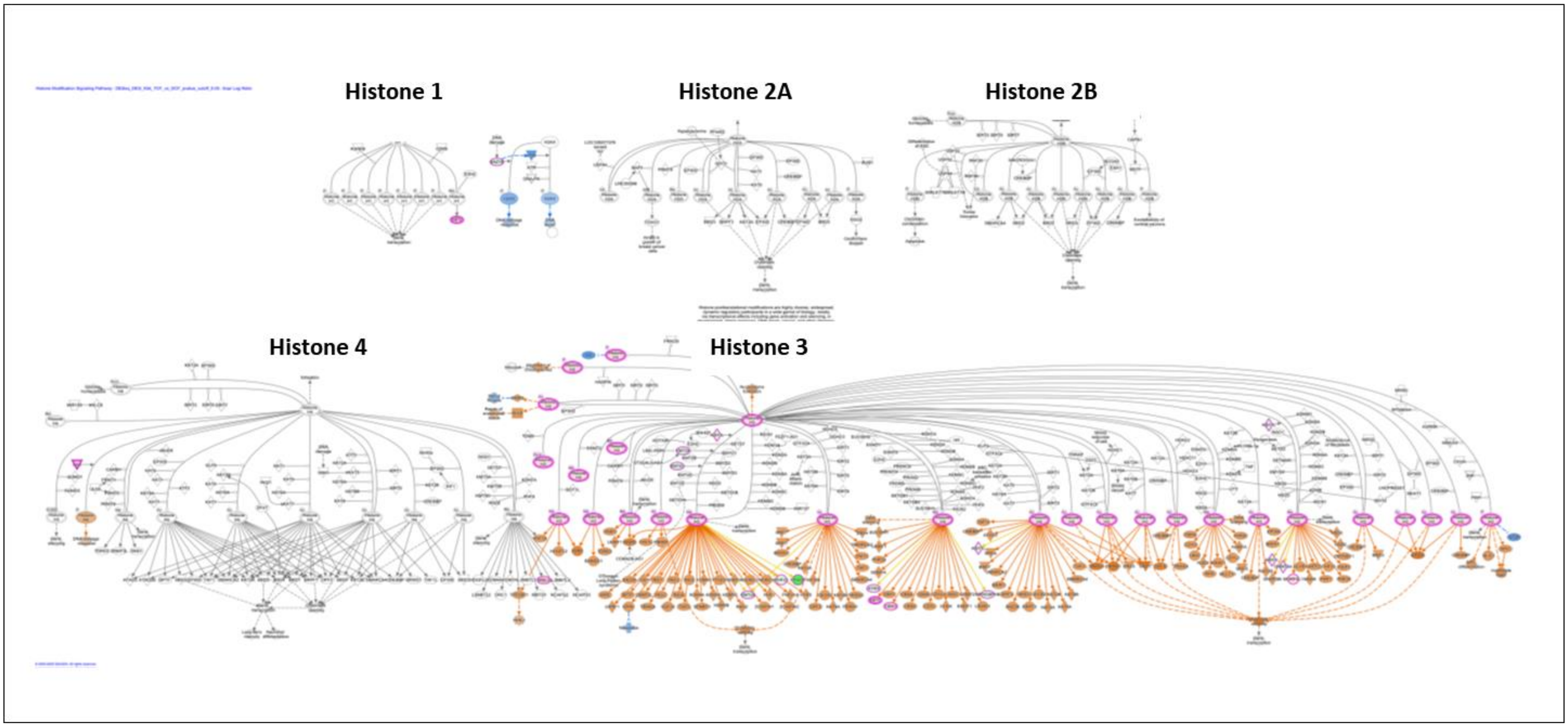


**Supplementary Fig S4**. Magnification of Fig. 4A.

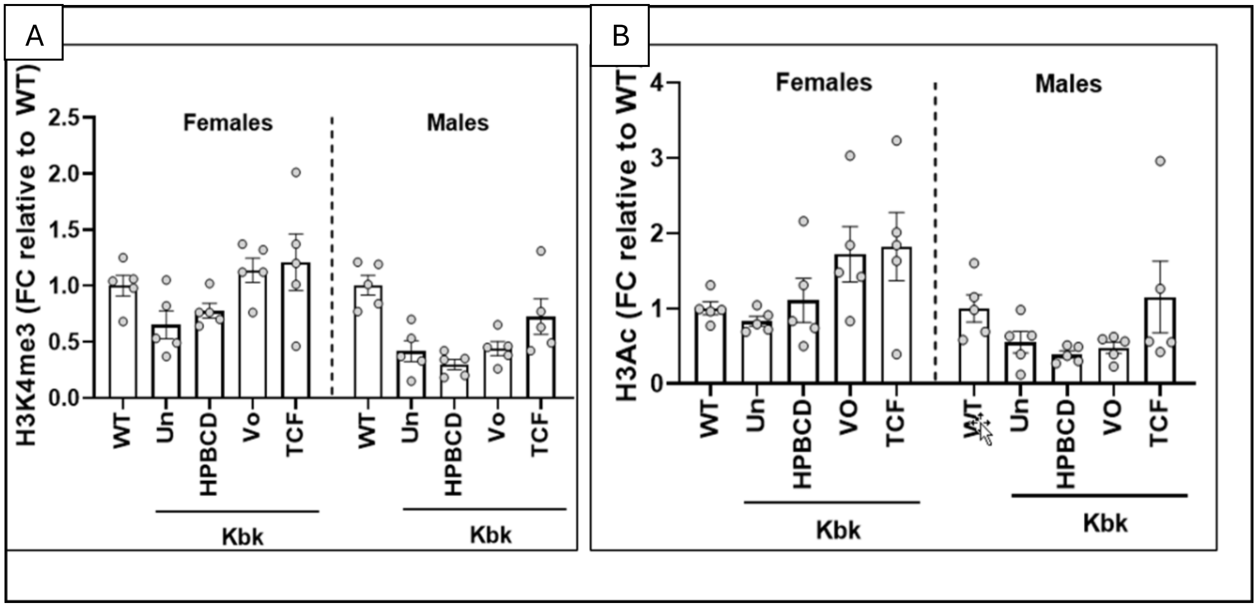


**Supplementary Fig. S5. Effects of sex on TCF-induced increase in histone H3K4 trimethylation and acetylation in mouse brain. A-B**. Quantitative analysis of western blots showing fold change (FC) of H3K4me3 and histone 3 acetylation (H3Ac) in the brain after the treatments. Kbk mice were given TCF or its components as indicated, starting at P21-P23, and the brain was analyzed at 3 months of age. WT mice were untreated. All data were normalized for total histone protein and fold change is relative to the average of WT mice. Each group consisted of 5 males or 5 females. Data are mean±SEM.

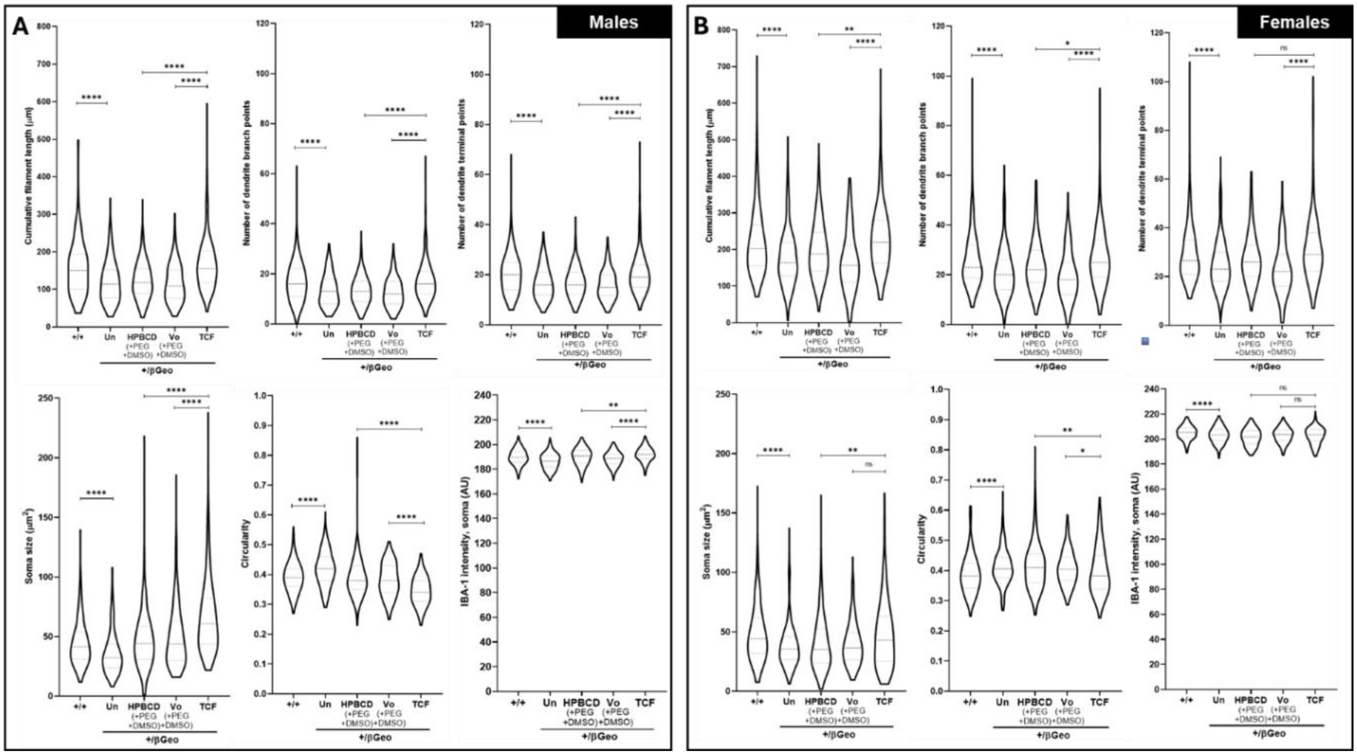


**Supplementary Fig. S6**. Quantitative morphometric analysis of hippocampal microglia in male and female Kbk mice. **A-B**. Violin plot showing quantitative results of microglia filaments (length, branch and terminal points) and soma (size, circularity and IBA1 intensity) in untreated and treated (A) males and (B) females. Treatments were as indicated. Approximately 200-350 microglia (~30-75 per mouse) from each gender were analyzed. Microglia were imaged in two random fields at two consistent depths of the hippocampus in each mouse. Males, n=4; females, n=4-5, analyzed at 3 months of age. WT, wild type; Un, untreated; TCF, triple combination formulation. Statistical analysis by Kruskal-Wallis test. ns, non-significant. **P<0.05, **P<0.01, ****P<0.0001.*

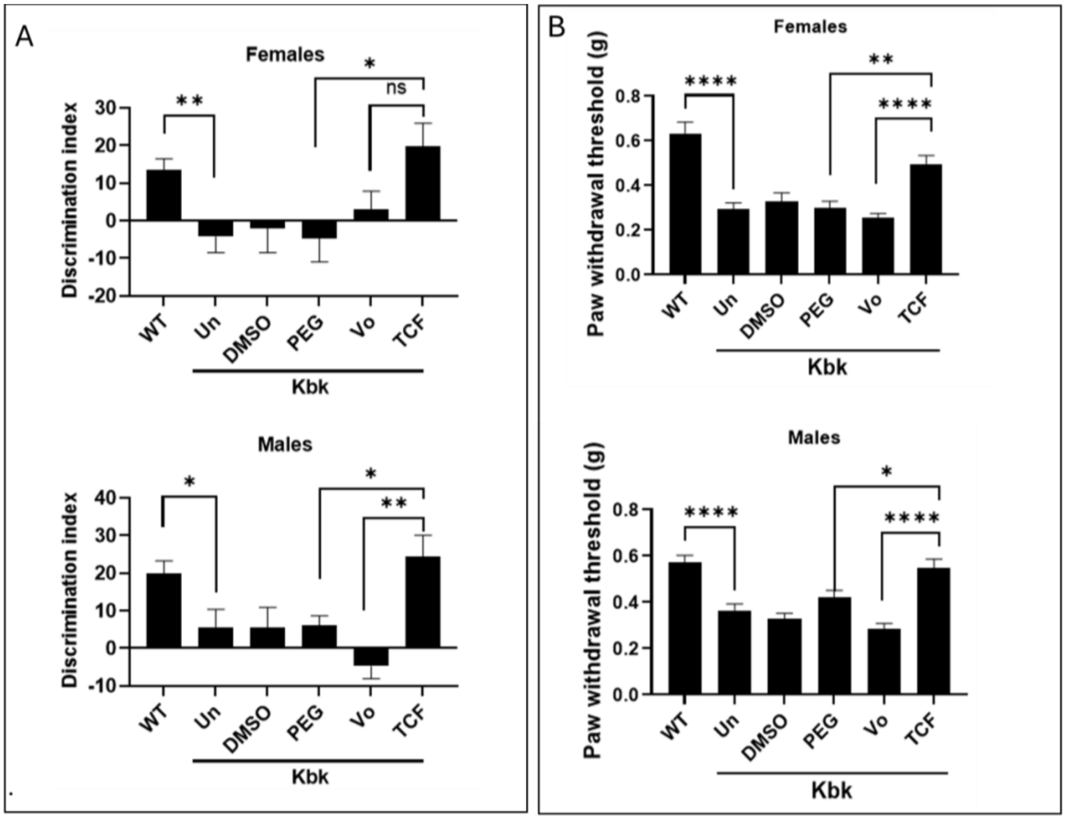

**Supplementary Figure S7. Sex-based evaluation of memory and tactile responses in drug-treated Kbk mice (A)** Bar diagrams show discrimination index determined by novel object recognition assay in females (upper panel) and males (lower panel) **(B)** Paw withdrawal threshold, an indicator of tactile sensitivity assessed by the Von Frey assay in female (upper panel) and male (lower panels) mice. Kbk animals were left untreated (Un) or given weekly i.p injections of TCF or its components as indicated starting at P21-P23 and assessed at 5-7 months of age. WT mice served as healthy control. Number of mice are as follows: WT males (n=13), females (n=12); untreated Kbk males (n=14), females (n=10); DMSO males (n=5), females (n=5F); PEG in DMSO, males (n=6), females (n=6); HPBCD in PEG and DMSO, males (n=9), females (n=11); and TCF males (n=9), females (n=9). Data are mean±SEM. Statistical analysis by Student's t-test. ns, non-significant. **P<0.05, **P<0.01, ***P<0.0001.*
